# A quantitative framework reveals the ecological drivers of grassland soil microbial community assembly in response to warming

**DOI:** 10.1101/2020.02.22.960872

**Authors:** Daliang Ning, Mengting Yuan, Linwei Wu, Ya Zhang, Xue Guo, Xishu Zhou, Yunfeng Yang, Adam P. Arkin, Mary K. Firestone, Jizhong Zhou

## Abstract

Unraveling the drivers controlling community assembly is a central issue in ecology. Selection, dispersal, diversification and drift are conceptually accepted as major community assembly processes. Defining their relative importance in governing biodiversity is compellingly needed, but very challenging. Here, we present a novel framework to quantitatively **i**nfer **c**ommunity **a**ssembly **m**echanisms by **p**hylogenetic bin-based null model analysis (iCAMP). Our results with simulated microbial communities showed that iCAMP had high accuracy (0.93 - 0.99), precision (0.80 - 0.94), sensitivity (0.82 - 0.94), and specificity (0.95 - 0.98), which were 10-160% higher than those from the entire community-based approach. Applying it to grassland microbial communities in response to experimental warming, our analysis showed that homogeneous selection (38%) and “drift” (59%) played dominant roles in controlling grassland soil microbial community assembly. Interestingly, warming enhanced homogeneous selection, but decreased “drift” over time. Warming-enhanced selection was primarily imposed on Bacillales in Firmicutes, which were strengthened by increased drought and reduced plant productivity. This general framework should also be useful for plant and animal ecology.

## Introduction

Microorganisms are the most diverse group of life presently known, inhabiting almost every imaginable environment on the Earth, and they typically form complex communities whose structure, functions, interactions, and dynamics are critical to our society. However, characterizing such complex communities, quantifying the accompanying ecological processes, and dissecting the underlying mechanisms are extremely challenging^1^. With the recent development of high-throughput metagenomic technologies^1^, large experimental data on community structure can be rapidly obtained. However, analyzing such massive data to address fundamental ecological questions such as community assembly mechanisms is challenging.

Understanding community assembly rules is a longstanding issue of ecologists^2-5^. While niche-based theory asserts that deterministic processes largely control the patterns of species composition, abundance and distributions, neutral theory assumes that all species are ecologically equivalent, and species dynamics is controlled by stochastic processes of birth/death, speciation/extinction, and immigration^6,7^. After intensive debates on niche vs neutral processes since 2000s^7-9^, it is generally accepted that both deterministic and stochastic processes operate simultaneously in the assembly of local communities^9-11^, and the key question becomes how to define their relative importance in controlling community diversity, distribution and succession^5,9-16^.

Under the deterministic–stochastic umbrella, Vellend^13,17^ proposed a conceptual framework that community diversity and dynamics are controlled by four high-level general ecological processes: selection, dispersal, speciation or diversification, and ecological drift^17,18^. Hereafter, we use the term “ecological processes” particularly to represent these community assembly processes. Although the framework has recently received a great attention in microbial ecology^5,19-21^, translating this conceptual framework into a quantitative operational model is even more challenging^18-20,22^. Most analyses with respect to the relative importance of the four processes across different types of natural communities are qualitative and subjective, and replete with great uncertainty^18^. As an exploratory effort, a null modelling-based operational approach was developed to obtain quantitative information on community assembly processes from the statistical perspective^21,22^, which is abbreviated as QPEN (Quantifying assembly **P**rocesses based on **E**ntire-community **N**ull model analysis) hereafter. QPEN uses phylogenetic metrics to infer selection since phylogenetic distance could reflect niche difference (so called phylogenetic signal) within some threshold^21,23^. Based on QPEN, selection under homogeneous abiotic and biotic conditions in space and time is referred to as constant selection^18^ or homogeneous selection (HoS)^22^, by which low phylogenetic compositional variations or turnovers are expected. In contrast, selection under heterogeneous conditions leads to high phylogenetic compositional variations, which is referred to as variable selection^18,22^ or heterogeneous selection (HeS)^5^. Similarly, dispersal is also divided into two categories^21,22^ - homogenizing dispersal (HD) and dispersal limitation (DL). The former refers to the situation that high dispersal rate can homogenize communities and hence lead to little taxonomic compositional variations, whereas the later signifies the circumstance that low dispersal rates could increase community taxonomic variations. When neither selection nor dispersal is dominated, community assembly is governed by drift, diversification, weak selection and/or weak dispersal, which is referred to be “undominated”^22^. For convenience, hereafter, the “undominated” is simply designated as “drift” (DR)^21^.

This statistical approach represents a significant advance in microbial ecology because it was the first time that microbial ecologists are able to obtain quantitative information on community assembly processes^5^. It has provided valuable insights into the importance of various ecological processes in microbial ecology^21,22,24-30^ and plant ecology^31^. However, one major limitation is that various ecological processes are quantified based on the pairwise turnovers of entire communities^21,22^ rather than the much finer biological levels on which ecological processes actually act, such as genes, genotypes, cells, individuals, populations, and lineages^11,19,20,32,33^. Within a single microbial community, certain populations are under strong selection, whereas others could be under strong drift. This type of difference cannot be discerned using whole community level metrics. Also, the responses to the changes in environmental conditions vary greatly among different groups of organisms. Similarly, the dispersal ability, diversification rates and susceptibility to drift are substantially different among various microbial groups. Thus, it would be meaningful to consider selection and other ecological processes at the level of individual taxa/lineages rather than the entire community^5,19^. To this end, we developed a general framework to quantitatively infer Community Assembly Mechanisms by Phylogenetic-bin-based null model analysis, abbreviated as iCAMP. We applied this novel approach to investigate whether and how experimental warming affects various ecological processes in the assembly of grassland soil microbial communities. Our results indicated that iCAMP provides a robust, reliable tool for quantifying the relative importance of ecological processes in controlling microbial community diversity and succession.

## Results

### Overview of iCAMP

To quantify various ecological processes, the observed taxa are first divided into different groups (“bins”) based on their phylogenetic relationships (Fig. 1, Fig. S1a). Then, the process governing each bin is identified based on null model analysis of the phylogenetic diversity using beta Net Relatedness Index (βNRI), and taxonomic β-diversities using modified Raup-Crick metric (RC) (Fig. 1, S1b). For each bin, the fraction of pairwise comparisons with βNRI < −1.96 is considered as the percentages of HoS, whereas those with βNRI > +1.96 as the percentages of HeS. Next, taxonomic diversity metric RC is used to partition the remaining pairwise comparisons with |βNRI| ≤ 1.96. The fraction of pairwise comparisons with RC ≤ −0.95 is treated as the percentages of HD, while those with RC > + 0.95 as DL, and the remains with |βNRI| ≤ 1.96 and |RC| ≤ 0.95 as the percentages of DR (Fig. 1). The above analysis is repeated for every bin. Subsequently, the fractions of individual processes across all bins are weighted by the relative abundance of each bin, and summarized to estimate the relative importance of individual processes at the whole community level (Fig. 1, S1e). Furthermore, various statistical analyses (e.g. correlation, regression, Mantel test, variation partitioning) are used to reveal the linkages of individual processes to different environmental factors for obtaining detailed insights into community assembly mechanisms (Fig. 1).

**Fig. 1.**
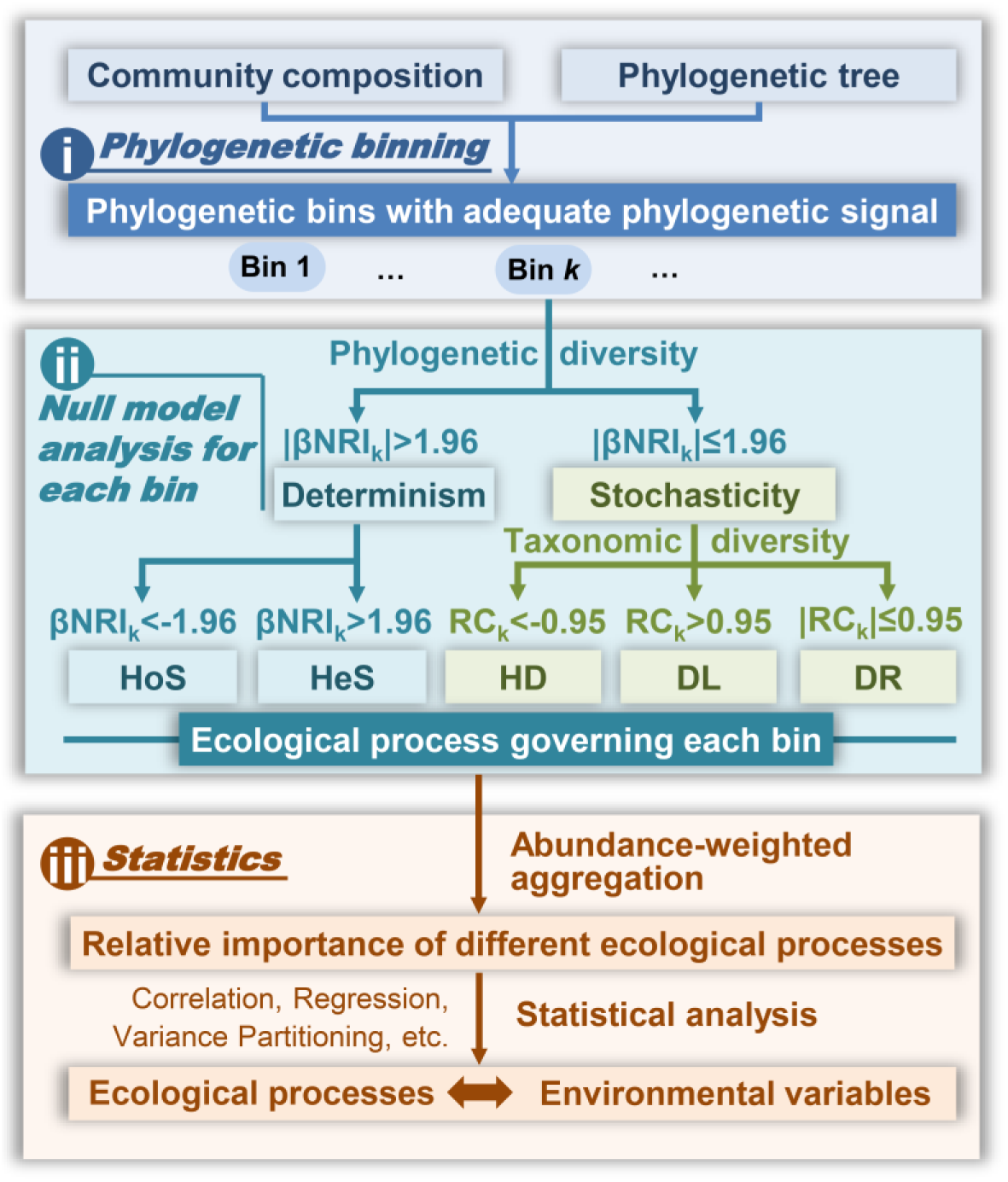
Overview of iCAMP. iCAMP includes several key steps: (i) phylogenetic binning, (ii) bin-based null model simulations with phylogenetic diversity for partitioning selection, and taxonomic diversity for partitioning dispersal and drift, and (iii) statistical analysis for assessing relative importance of different ecological processes and linking the processes with different environmental factors. βNRI, beta net relatedness index; RC, modified Raup-Crick metric; Here, “ecological processes” particularly mean community assembly processes, including homogeneous selection (HoS), heterogeneous selection (HeS), homogenizing dispersal (HD), dispersal limitation (DL), and “drift” (DR). See main text for detailed explanation.

### Simulated communities

Due to lack of a gold-standard experiment to establish the true community assembly processes, using simulated communities is the predominant approach for assessing performances of various computational methods, as shown in many other computational studies^34-36^. Thus, we used a simulation model to generate communities with pre-defined relative importance of each ecological process (Fig. S2). In this simulation model, four plots (LA, LB, HA, HB) in two islands (A, B) under two types of environments (L, H) are considered, with six local communities sampled from each plot (Fig. S2a). Each local community consists of different types of species controlled by drift, selection, or dispersal (Fig. S2c, d, e).

Three scenarios were simulated with three different levels of phylogenetic signal in the regional species pool: low (Blomberg’s K = 0.15), medium (K = 0.9), and high (K = 5.5). Each scenario has 15 simulated situations, where the expected (“true”) relative importance of each process (selection, dispersal, drift) was set from 0% to 100% (with an interval of 25%, Fig. S2b and Table S1). Then, the relative importance of different ecological processes were assessed by iCAMP. The performance was evaluated with six quantitative and qualitative indexes: quantitative accuracy and precision, and qualitative accuracy, precision, sensitivity, and specificity (detailed in Methods, Eqs. 16-21). The quantitative performance indexes are based on the difference between the expected and estimated relative importance of each process, while the qualitative performance indexes are calculated from the true or false identification of dominant process.

According to the performances with simulated communities, we optimized various algorithms and parameter settings for iCAMP (detailed in Supplementary Text A). Our results revealed that: (i) Tree-based phylogenetic binning (Fig. S3c) was preferred compared to the other two algorithms based on phylogenetic distances (Fig. S3a, b, d, e); (ii) The optimized minimal requirement of bin size (*n*_*min*_ = 24 species) led to optimal within-bin phylogenetic signal (Fig. S4); (iii) βNRI had better performance than βNTI (beta Nearest Taxon Index) in iCAMP (Fig. S5); (iv) The phylogenetic null model for βNRI showed better performance when randomizing taxa within bin, while taxonomic null model for RC performed better using across-bin randomization (Fig. S6); (v) Randomization for 1,000 times in null model analysis should be adequate in terms of reproducibility (Fig. S7); and (vi) Reducing taxa number is inadvisable for iCAMP (Fig. S8).

### Comparison between iCAMP and other approaches

After appropriate parameter settings were determined, iCAMP and several previously reported approaches were compared for their performances with the simulated communities (Fig. 2, Fig S9, S10). First, the ecological stochasticity was quantified by five approaches, including abundance-weighted neutral taxa percentage (NP)^5^ based on neutral-theory model, normalized stochasticity ratios (NST)^37^ based on taxonomic (tNST) or phylogenetic metrics (pNST), and the relative importance of stochastic processes (HD, DL, and DR) based on QPEN or iCAMP. Under high- and medium-phylogenetic-signal scenarios, iCAMP always showed the highest quantitative accuracy (0.978 - 0.997) and precision (0.903 - 0.930), while pNST exhibited similar accuracy (0.924 - 0.954) but lower precision (0.658 - 0.722, *p* < 0.001, Fig. 2a-c). Under low-phylogenetic-signal scenario, iCAMP still had the highest precision (0.807) and the second-high accuracy (0.770), while pNST showed similar precision (0.723) and the highest accuracy (0.947). In contrast, tNST, NP, and QPEN showed significantly lower precision (<0.57, down to negative, *p* < 0.0001) than iCAMP across all scenarios (Fig. 2a-c).

**Fig. 2.**
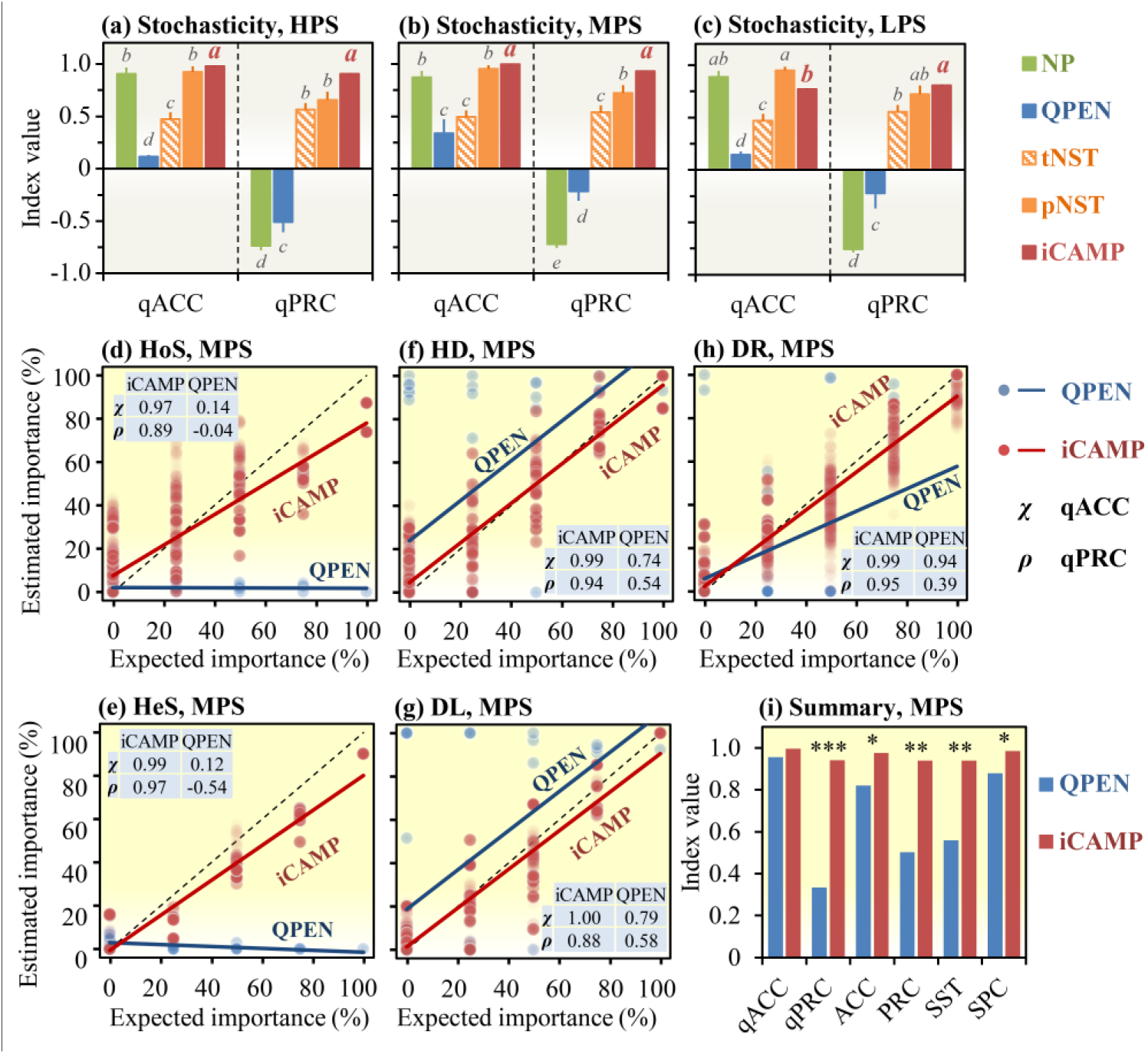
Performances of iCAMP and other approaches with simulated communities. **(a-c)** The performances of different approaches in quantifying ecological stochasticity under high- (HPS), medium- (MPS), and low-phylogenetic-signal (LPS) scenarios, which were assessed by quantitative accuracy (qACC) and precision (qPRC). (**d-h**) The performances of iCAMP and QPEN were also evaluated based on the consistency between the estimated and expected relative importance of different individual ecological processes. **(i)** Overall performance of iCAMP and QPEN under MPS scenario assessed with six performance indexes: qACC (*χ*), qPRC (*ρ*), qualitative accuracy (ACC), qualitative precision (PRC), sensitivity (SST), and specificity (SPC). NP, abundance-weighted neutral taxa percentage; tNST and pNST, taxonomic and phylogenetic normalized stochasticity ratio. Significance was indicated as ***, *p* < 0.001; **, *p* < 0.01; *, *p* < 0.05; or by different letters (*p* < 0.05).

Since only QPEN and iCAMP can quantify relative importance of different ecological processes, their performances were particularly compared. Overall, iCAMP had, on average, higher accuracy (9.9% higher), precision (120.2%), sensitivity (61.1%), and specificity (10.6%) than QPEN (Fig. 2i, S9k, l). The setting of phylogenetic signal also had significant impacts on iCAMP performance. When the phylogenetic signal increased from low/medium (Fig. 2i, S9k) to high (Fig. S9l), the accuracy and specificity of iCAMP remained high (> 0.92) without any significant change (*p* > 0.20), but the precision and sensitivity of iCAMP increased from 0.80-0.82 to 0.90-0.94. In contrast, the overall performance of QPEN did not show improvement when phylogenetic signal increased.

The performance varied considerably among different ecological processes (Fig. 2d-h, S10). In the simulated communities, QPEN had lower performance (quantitative precision < 0, qualitative precision < 0.13, sensitivity < 0.04, Fig. S10) in estimating HoS and HeS even under high phylogenetic signal (Fig. S9f, g). iCAMP improved the estimation of all processes. Under medium and high phylogenetic signals, all performance indices were higher than 0.78 for iCAMP (Fig. S10). However, with low phylogenetic signal, iCAMP had low sensitivity (down to 0.17) for HoS (Fig. S10), despite that it was considerably (*p* < 0.05) higher than QPEN (sensitivity < 0.04 for HoS). These results confirmed that low phylogenetic signal of niche preference can limit the capability of phylogenetic metrics to infer selection, which can be partly but not completely overcome by iCAMP. Nevertheless, the quantitative performance of iCAMP remained relatively high for all processes under all scenarios (quantitative accuracy and precision > 0.71). Collectively, all these results indicated that iCAMP can substantially improve the quantitative estimation of community assembly processes.

### Effects of warming on bacterial assembly processes in a temperate grassland

To determine the effectiveness of iCAMP in real-world studies, iCAMP was applied to an empirical data of soil bacterial communities in a grassland under experimental warming^38^. Based on iCAMP analysis, HoS and DR were more important than other processes in bacterial community assembly, with average relative importance of 37.0-38.5% and 58.3-59.9% (Fig. 3a, b), respectively. Warming significantly altered the relative importance of different processes (*p* < 0.01, permutational ANOVA). Since other processes had quite low estimated relative importance (< 3.4%), we primarily focused on the effects of warming on HoS and DR in subsequent analyses. Overall, warming decreased the relative importance of DR and increased HoS. Significant year-to-year variations were observed (Fig. 3c, d). In the first year, the communities under warming showed significantly higher ratio of DR (Cohen’s *d* = 2.9, *p* = 0.001), but lower ratio of HoS (Cohen’s *d* = −2.7, *p* < 0.001) than those under control, suggesting the bacterial community assembly was even more stochastic under warming than control in the beginning. In the second year, the difference between warming and control became insignificant. In the third to fifth years, the communities under warming had significantly higher ratio of HoS (Cohen’s *d* = 0.6-1.7) and lower ratio of DR (Cohen’s d = −0.8 to −1.3), suggesting that the selection pressure imposed by warming on the soil bacteria gradually increased with time.

**Fig. 3.**
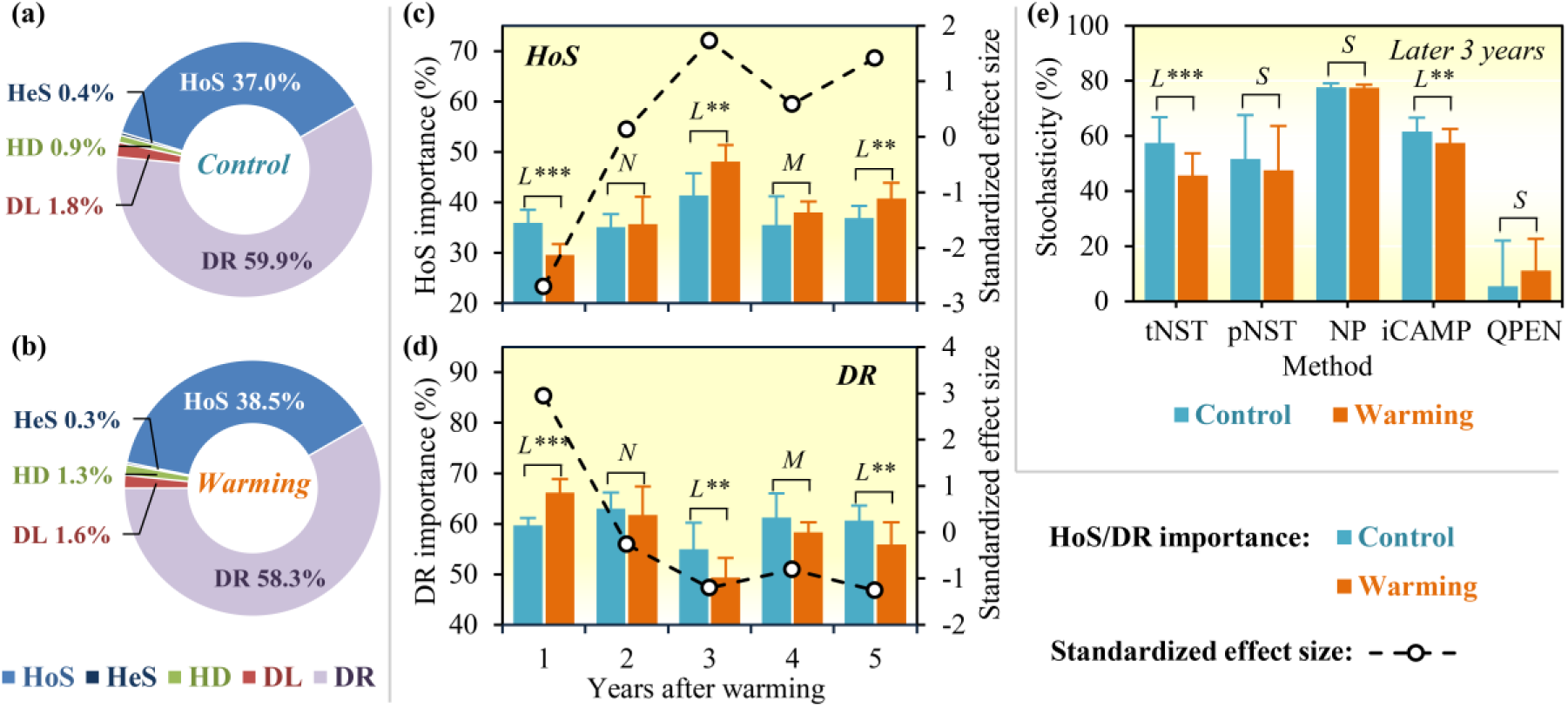
Relative importance of different ecological processes estimated with iCAMP in response to experimental warming. **(a)** Under Control. (**b**) Under warming. **(c)** Changes of HoS under warming (orange bar) and control (aqua bar), and the effect size of warming (black dashed line) on HoS across different years after warming. **(d)** Changes of DR under warming and control and the effect size of warming on DR. **(e)** Stochasticity estimated by different methods in the later 3 years. The standardized effect size of warming was Cohen’s *d*, estimated as the mean difference between warming and control divided by pooled standard deviation. Error bars represented standard deviations. Significance was expressed as ***, *p* < 0.01; **, *p* <0.05; L, M, S, and N represented large, medium, small, and negligible effect sizes.

QPEN was also applied to quantify the ecological processes. The results from QPEN indicated that HoS predominated (> 73%) bacterial assembly with higher relative importance under warming (83.3%, Fig. S11a) than control (73.3%, Fig. S11b) although not significant (*p* = 0.174). QPEN suggested 0.0% of DR under warming, 0.0 % of HeS and HD across all years, and 100% of HoS in some years (Fig. S11c, d).

### Stochastic vs deterministic bacterial assembly in the grassland

In microbial ecology, DL, HD, and DR are generally considered stochastic (neutral) processes^5^, thus the sum of their estimated relative importance can be used to estimate stochasticity of community assembly. Based on iCAMP results, the relative importance of stochastic processes was 62.6% under control and 61.3% under warming (Fig. S12). In contrast, QPEN estimated the relative importance of stochastic processes was 26.7% under control and 16.7% under warming, which were much lower than those estimated by other approaches (Fig. S12). For instance, variation partitioning analysis (VPA) revealed that substantial portions of the community variations (68.4%) could not be explained by all measured environmental variables^38^. The tNST and pNST were on average 48.8% under warming and 52.3% under control, and NP ranged from 74% to 79% in different years for both warming and control (Fig. S12). Obviously, VPA, NST, and NP showed more consistent results with iCAMP than QPEN.

All approaches did not reveal significant (*p* > 0.10) differences of the 5-year mean stochasticity between warming and control, except tNST with medium effect size (*p* < 0.05, Fig. S12). But in the third to fifth year (Fig. 3e), both tNST and iCAMP revealed that warming had significant (*p* < 0.05) decrease in stochasticity, and there was slight decrease in stochasticity with pNST and NP under warming (small effect size) though it was insignificant (*p* > 0.10). On the contrary, QPEN showed a slight but insignificant increase in stochasticity. Collectively, consistent with our previous analysis^38^, various approaches except QPEN supported that stochastic processes could play more important roles in grassland soil bacterial assembly and that warming decreased the stochasticity after three years.

### Variations of assembly mechanisms across different phylogenetic groups

Distinct from previous approaches, iCAMP can provide information on the relative importance of different ecological processes in individual lineages or bins. For this purpose, the observed 18,123 OTUs were divided into 658 phylogenetic bins, each of which was then analyzed separately as outlined in Fig. 1. Our results revealed that HoS dominated 59 bins (9% of bin numbers and 33% of relative abundance, Fig. 4a). Two of the major bins were Bacillales (Bin 1, 26.7% in total abundance of HoS-controlled bins) in Firmicutes and Spartobacteria (Bin 2, 18.8%) in Verrucomicrobia (Fig. 4b, S13a). In contrast, DR dominated 598 bins (91% of bin numbers and 67% of relative abundance, Fig. 4a), which mainly belonged to Class Alphaproteobacteria (22.2% in total abundance of DR-controlled bins) and Phylum Actinobacteria (23.5%) (Fig. S13a).

**Fig. 4.**
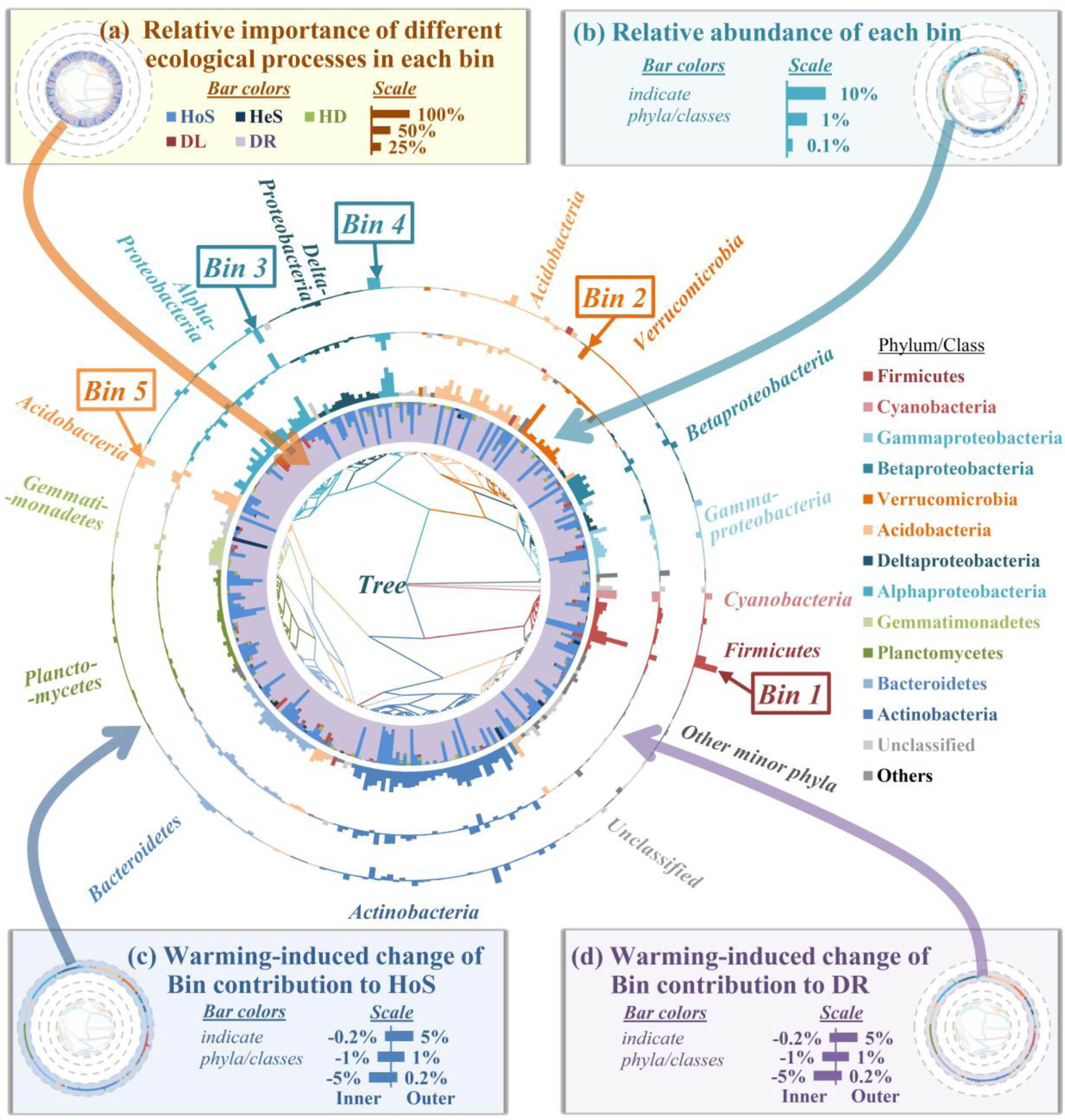
Variations of ecological processes across different phylogenetic groups. Phylogenetic tree was displayed at the center. **(a)** Relative importance of different ecological processes in each bin (stacked bars in 1^st^ annulus), HoS, blue bar, HeS, dark blue bar, HD, green bar, DL, red bar, and DR, violet bar; **(b)** Relative abundance of each bin (2^rd^ annulus); **(c)** Warming-induced change in bin contribution to HoS (4^th^ annulus); and **(d)** DR (3^th^ annulus) in later 3 years, where positive (outward bar) and negative (inward bar) represented increase and decrease by warming, respectively. Only the top 300 bins were shown in this figure, accounting for a total relative abundance of 95%. Bin 1 to Bin 5 were the five most abundant bins.

To understand how different lineages respond to warming, we further determined the bacterial groups contributing to the warming-induced changes of HoS and DR in the third to fifth years (Fig. 4c, d). Our results revealed that Firmicutes contributed 58.2% of the warming-induced increases in HoS (Fig. S13b). The most abundant Firmicutes bin (Bin 1, Bacillales, average 74.8% in Firmicutes) was always governed by HoS (Fig. S13c). Warming gradually drove its abundance significantly (*p* < 0.05) higher than those under controls (Fig. S13c). In contrast, the decrease of DR under warming was due to similar negative responses of many bins in five phyla (Proteobacteria, Verrucomicrobia, Bacteroidetes, Planctomycetes, and Acidobacteria, Fig. 4c, S13b). For instance, Bin 4 of Rhizobiales in Alphaproteobacteria (Bin 4. Fig. S13b) had lower relative abundance and reduced relative importance of DR under warming, especially in later 3 years (Fig. S13d). These results demonstrated that different bins were controlled by quite distinct assembly mechanisms in response to warming.

### Environmental factors influencing ecological processes

Plant and environmental factors also affected the relative importance of different ecological processes (Fig. 5, S14, S15, Supplementary Text B). Overall, the two major processes (HoS and DR) were significantly correlated with the factors related to precipitation, plants, soil nitrogen, and temperature (Fig. 5). All these factors showed distinct association strengths with HoS or DR under warming and control.

**Fig. 5.**
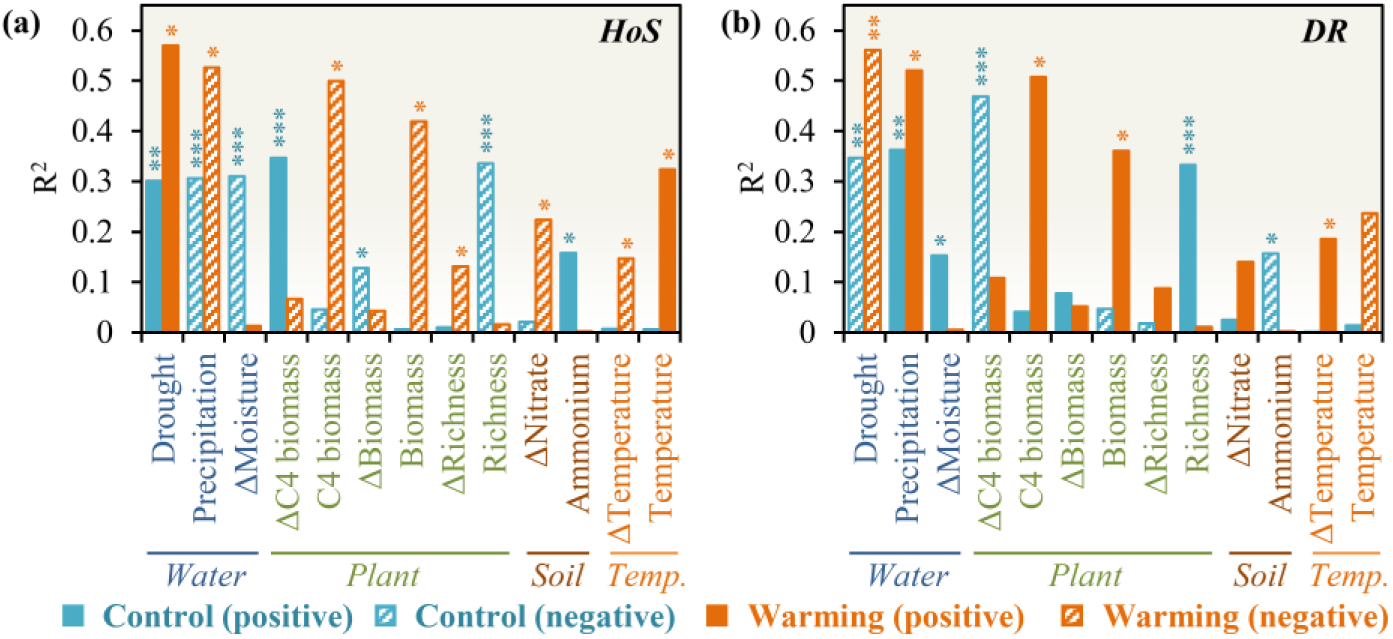
Effects of environmental factors on the major ecological processes under warming (orange) and control (aqua). **(a)** Correlations of individual environmental factors with HoS, and **(b)** “Drift” (DR). R^2^, coefficient of determination based on the best model from mcMantel analysis (see Table S2 for details). The correlation was determined based on the difference (marked with “Δ”) or the mean (without “Δ”) of a factor between each pair of samples. In this figure, drought, precipitation, plant, soil temperature and other properties were measured in the sampling month, while moisture values were annual means. Only factors significantly correlated with HoS or DR were shown, see Table S2 for other factors. Significance was expressed as ***, *p* < 0.01; **, *p* < 0.05; *, *p* < 0.1.

The precipitation and plant-related factors showed stronger association with HoS and DR than all other factors. The precipitation and drought index in the sampling month showed significant correlation with HoS (Fig. 5a, S14a) and DR (Fig. 5b), with slightly stronger association under warming (R^2^ = 0.52-0.57, *p* < 0.1, Fig. 5a, b) than control (R^2^ = 0.30-0.36, *p* < 0.05, Fig. 5a, b). This suggests enhanced selection pressure when warming and drought occurred simultaneously in low-precipitation years. Plant biomass showed strong correlation with HoS and DR under warming (Fig. S14b; R^2^ > 0.36, *p* < 0.1, Fig. 5), but not under control (R^2^ < 0.05, *p* > 0.4). In contrast, the plant richness and variation of C-4 plant biomass significantly correlated with HoS and DR under control (Mantel R^2^ > 0.33, *p* < 0.01, Fig. 5), but not under warming (R^2^ < 0.1, *p* > 0.3). Soil temperature in the sampling month was significantly correlated with HoS under warming (Mantel R^2^ = 0.32, *p* < 0.1, Fig. 5a; 2^nd^ rank in MRM, Fig. S15b), but not under control (R^2^ < 0.01, *p* > 0.7), suggesting that warming enhanced the filtering effects of soil temperature.

## Discussion

Disentangling ecological drivers controlling community assembly is crucial but difficult in ecology, especially in microbial ecology. Although metagenomics and associated technologies have revolutionized microbial ecology research^1^, a great challenge is how to use such massive data to address compelling ecological questions such as community assembly mechanisms. Thus, in this study, we described a novel framework, iCAMP, to quantify the relative importance of different ecological processes underlying community diversity and dynamics based on individual phylogenetic groups (bins) rather than the entire community. Various analyses demonstrated that iCAMP improved performance substantially with higher precision, sensitivity, specificity, and accuracy compared with previous approaches. The developed framework would help to move microbial ecology beyond biodiversity patterns towards mechanistic understanding of community diversity and succession. Although this framework is tested with microbial community data, it should also be applicable to plant and animal communities.

To quantify selection, phylogeny-based approaches^22-25,28,29,39-41^ require that the phylogenetic distances among taxa reflect their niche difference, i.e. there is phylogenetic signal or niche conservatism^5,23^. Although phylogenetic niche conservatism of microbial traits was reported^42,43^, the signals were mostly at medium or low levels (i.e. close to or lower than Brownian Motion expectation)^44^. Fortunately, significant phylogenetic signals were frequently found within a short phylogenetic distance ^29,40,42^, which was employed by recent microbial studies^24,28,29,41^, particularly in QPEN^21,22^. However, QPEN did not perform well in inferring selection with the simulated communities, possibly because it does not distinguish the differential influences of selection on distinct phylogenetic groups. In contrast, iCAMP partitions selection based on individual phylogenetic groups within a short phylogenetic distance, and hence it can greatly improve quantitative performance with accuracy and precision > 0.71 even when the overall (across-tree) phylogenetic signal is low.

A central question in ecology is to determine the relative importance of deterministic and stochastic processes in controlling community diversity and succession^5,9-16^. The results from iCAMP indicated that the grassland soil microbial community is more stochastic with an average of ∼60% stochasticity, while those from QPEN suggested that it is more deterministic with an average of 8% stochasticity, down to 0% in several years. We believe that the conclusion from iCAMP is more convincing due to two primary reasons. First, the iCAMP results are more consistent with several other commonly used methods such as multivariate analysis (e.g. VPA), null models (e.g. tNST, pNST), and neutral theory models (e.g. NP)^5^, which revealed an average of 48.8% to 79.0% stochasticity. Second, the results from iCAMP are more in accordance with our general expectation that stochastic processes should play more important roles under less stressful environments^10-12,14,16^. Since the grassland has rich C resources (above ground biomass, 272 g dry weight/m^2^ on average), intermediate level of precipitation (550 to 994 mm annual precipitation), mild temperature (∼15.6 °C mean annual air temperature), and nearly neutral soil pH (5.6 to 7.0), the grassland soil microbial communities appear to be not under very stressful conditions, and hence stochastic processes should play important roles, but at least not zero as revealed by QPEN.

Unraveling the drivers controlling the responses of ecological communities to climate change is also a critical topic in ecology and global change biology^38^. Several previous studies demonstrated that climate warming have significant impact on microbial diversity^45^, structure^38^, functional gene composition^46^, and activities^46-50^, but the underlying community assembly processes were rarely examined. In our grassland site, experimental warming increased the soil temperature by ∼3 °C^38^, thus it may gradually impose selective pressure as a deterministic force to decrease stochasticity as evident by our previous studies^38,45^. Here, iCAMP further revealed that warming gradually enhanced HoS and decreased DR in bacterial community assembly, and that the warming-induced selection was enhanced by drought and lower plant biomass. These results are consistent with several previous studies showing that the effects of warming on microbial communities intertwines with precipitation/drought^51-56^ and plant variables^57^. In addition, our results showed that the warming-enhanced HoS was mainly attributed to the positive responses of a group of Bacillales in Firmicutes, which are ubiquitous gram-positive bacteria in nature. Bacillales are endospore-forming and can remain in the dormant state for years^58^. These traits could offer their competitive advantages under selective pressure from increased temperature and drying. Finally, our experimental findings have an important implication for predicting and mitigating the ecological consequences of climate warming. Because warming enhances homogeneous selection, which leads community to be more similar, and decreases drift, the future community states and associated functions could be more predictable under warmed climate, and hence might be more manipulatable for mitigating climate change effects.

Although iCAMP has better performance over traditional approaches and provided valuable insights into the ecological processes governing the responses of grassland soil microbial communities to climate warming, there are still some limitations. For instance, one of the fundamental processes, diversification, is important to govern community assembly^17,19,20^, but it is not explicitly accounted for in iCAMP. Diversification is still embedded in DR with drift, weak selection, and/or weak dispersal in this framework. Environmental filtering and biotic interactions could also not be differentiated from each other in HoS and HeS. Thus, further developments are needed by incorporating null model of evolution to infer the relative importance of diversification, and by integrating functional traits-based network approaches to disentangle biotic interactions from abiotic filtering.

## Methods

### Procedure of iCAMP

iCAMP includes three major steps (Fig. 1). The first step is phylogenetic binning (Fig. S1a, S3). Three binning algorithms were compared. One is based on the distance to abundant taxa (Fig. S3a). The most abundant (i.e. the highest mean relative abundance in the regional pool) taxon is designated as the first bin to serve as the core taxon. All taxa with distances to the core taxon less than the phylogenetic signal threshold, *d*_*s*_, are assigned to this core taxon. The next bin is generated from the rest taxa in the same way. Consequently, a series of bins are generated with strict radiuses less than *d*_*s*_, so called strict bins. However, some strict bins may have too few taxa to provide enough statistical power for further analysis. Thus, each small bin is merged into its nearest neighbor bin until all bins reach the minimal size requirement, *n*_*min*_. The second algorithm is based on pairwise distances (Fig. S3b). The first bin consists the most abundant taxon, and all other taxa among which all pairwise distances are lower than *d*_*s*_. The second bin includes the next most abundant taxon among the remaining taxa. This procedure continues until all taxa are classified into different bins. To ensure each bin have enough size (≥ *n*_*min*_), a small bin less than *n*_*min*_ is merged into the nearest neighbor until all bins reach the minimal requirement *n*_*min*_. The third algorithm is based on phylogenetic tree (Fig. S3c). The phylogenetic tree is truncated at a certain phylogenetic distance (as short as necessary) to the root, by which all the rest connections between tips (taxa) are lower than the threshold *d*_*s*_. The taxa derived from the same ancestor after the truncating point are grouped to the same strict bin. Then, each small bin is merged into the bin with the nearest relatives. This procedure is repeated until all merged bins have enough taxa (≥ *n*_*min*_).

The objective of phylogenetic binning is to obtain adequate within-bin phylogenetic signal. To evaluate phylogenetic signal within each bin, the correlation between the pairwise phylogenetic distances and niche preference differences were analyzed by Mantel tests, where niche preference means the niche leading to optimum fitness (or relative fitness reflected by relative abundance) of a taxon. The bins with Pearson correlation coefficient R > 0.1 and *p* < 0.05 (one tail) are considered as bins with significant phylogenetic signal. In simulated communities, the niche preference difference between two taxa is treated as the key trait value difference. For empirical data, an index, i.e., niche value, is estimated as the relative-abundance-weighted mean of an environmental factor for each taxon as previously reported^23^. For instance, if OTU1 has relative abundances of 10%, 20%, and 10% in three samples under the temperature of 10, 20, and 30 °C, respectively, the temperature niche value of OTU1 is (10 × 10% + 20 × 20% + 3 × 10%) / (10% + 20% + 10%) = 20 °C. Then, the difference of niche values between taxa reflects niche difference, which are used for phylogenetic signal estimation. An optimized *n*_*min*_ should lead to the highest number of bins with significant phylogenetic signal and relatively high average correlation coefficient (average R) within bins. In this study, the optimized *n*_*min*_ is 24 for simulated datasets and 12 for the empirical data.

The second step is the null model analysis within each bin shown in Fig. S1b. Accordingly, an operator is defined as below to count whether a bin is governed by a process (Eq.1 to Eq. 10).

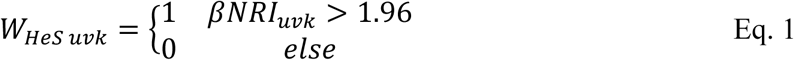

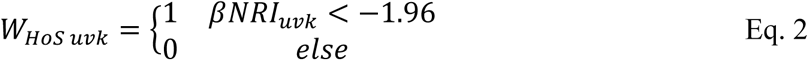

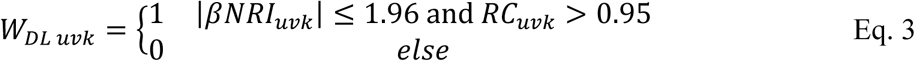

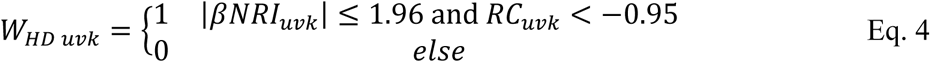

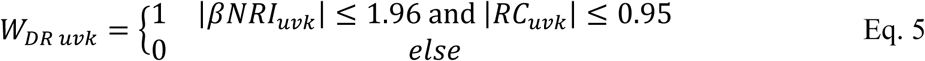

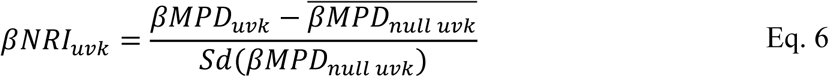

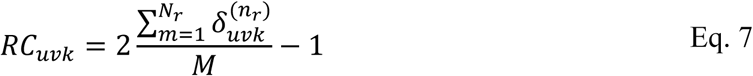

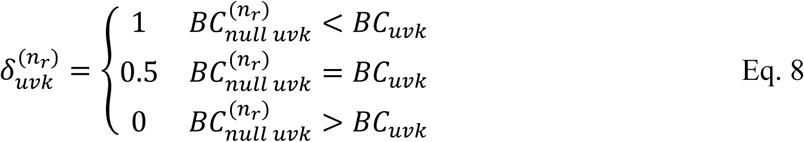

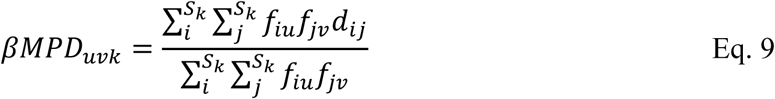

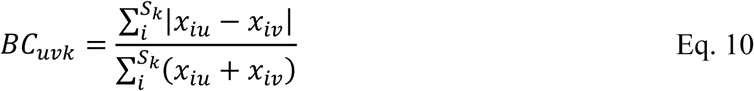

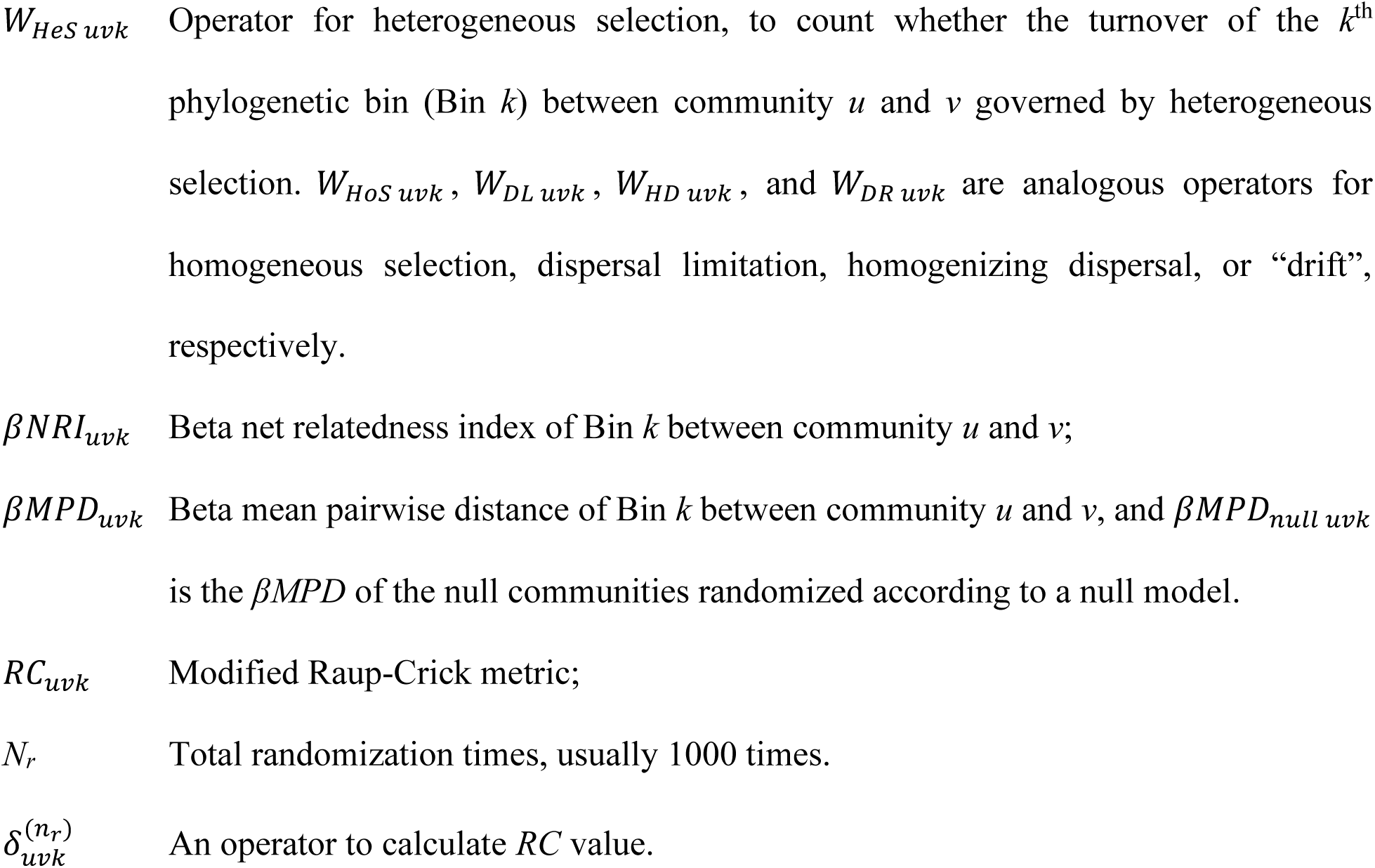
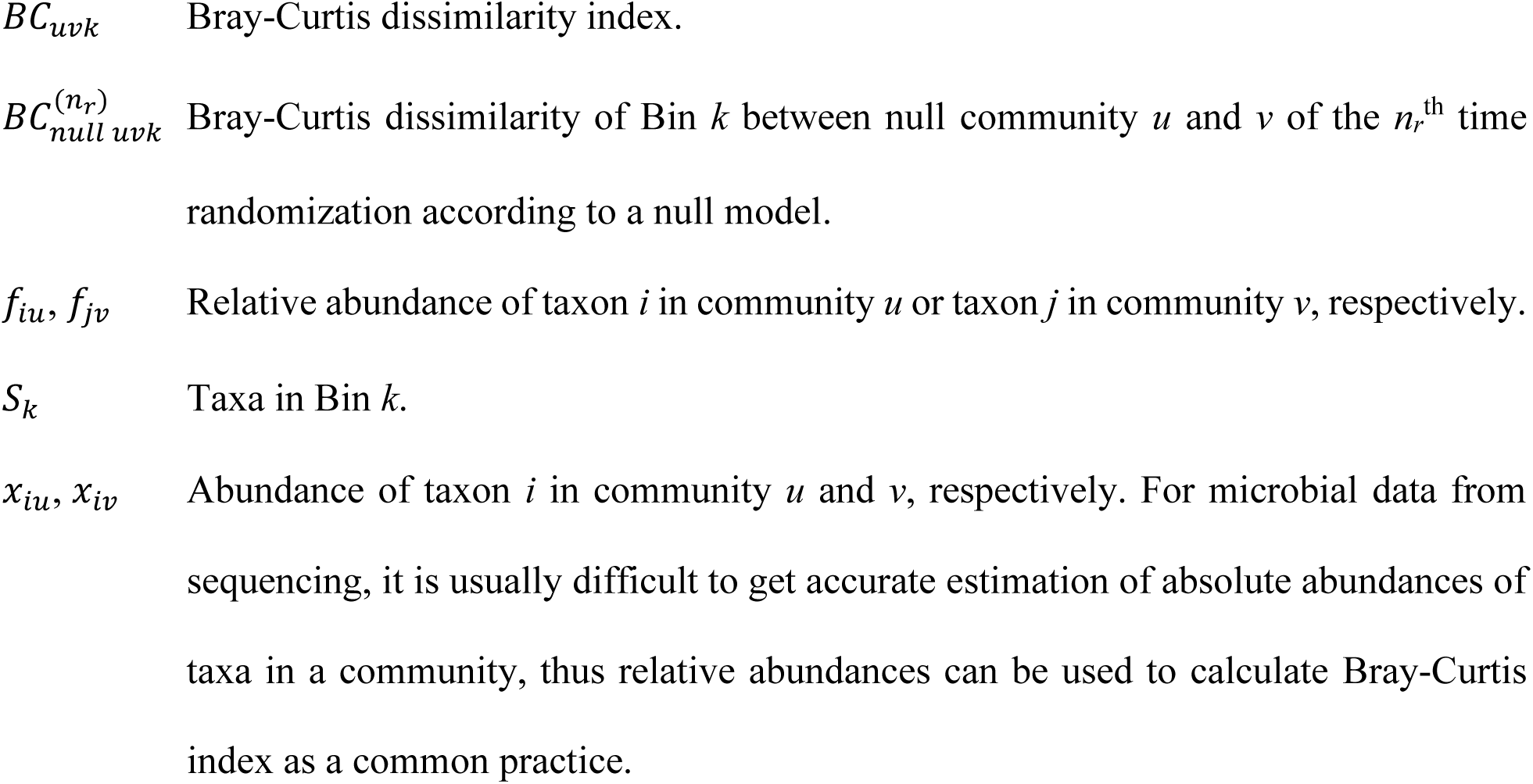

The null model algorithm for phylogenetic metrics is “taxa shuffle”^23,39^, which randomizes the taxa across the tips of the phylogenetic tree, and thus it randomizes the phylogenetic relationship among taxa. The null model algorithm for taxonomic metric is the one constraining occurrence frequency of each taxon proportional to observed and richness in each sample fixed to observed^21,59^. The null model algorithm results heavily depend on the selection of the regional pool, within which randomization is implemented^59^. Thus, the algorithms randomizing taxa within each bin and across all bins were compared in iCAMP analysis for the simulated communities.

Null model analysis is most computational resource- and time-consuming, which largely depends on the times of randomization and taxa number. But decreasing randomization times or taxa number can reduce reproducibility of the null model analysis. Considering that most reported null model analyses used 1,000-time randomization, iCAMP were performed for simulated data with randomization times ranging from 25 to 5,000 and repeated 12 times with each number of randomization times. The results from 60,000-time randomization served as a standard for evaluation. In addition, three methods for reducing taxa number were tested. The method “rarefaction” means to randomly draw the same number of individuals (sequences) from each sample and reduce the taxa number. The method “average abundance trimming” ranks all taxa from abundant to rare according to their average relative abundances across all samples and only keeps the taxa before a certain rank. The method “cumulative abundance trimming” ranks taxa in each sample from abundant to rare, then only keeps the abundant taxa in each sample so that every sample has the same cumulative abundance. The iCAMP results from the three methods were compared to that from the original simulated communities.

The third step of iCAMP is to integrate the results of different bins to assess the relative importance of each process (Fig. S1c-f). Defining neutrality at individual level has been proved a key to successfully develop the unified neutral theory^7^. Therefore, the relative importance of a process can be quantitatively measured as abundance-weighted percentage for each bin (Eq. 11) or the entire communities (Eq. 12Eq. 12 and 13). Qualitatively, for each pairwise comparison between communities (samples), the process with higher relative importance than other processes is regarded as the dominant process.

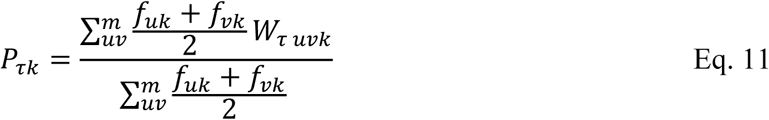

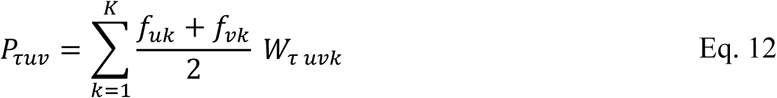

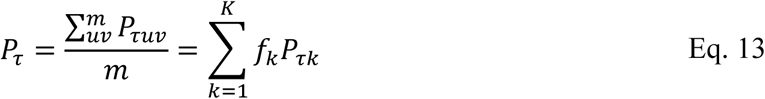

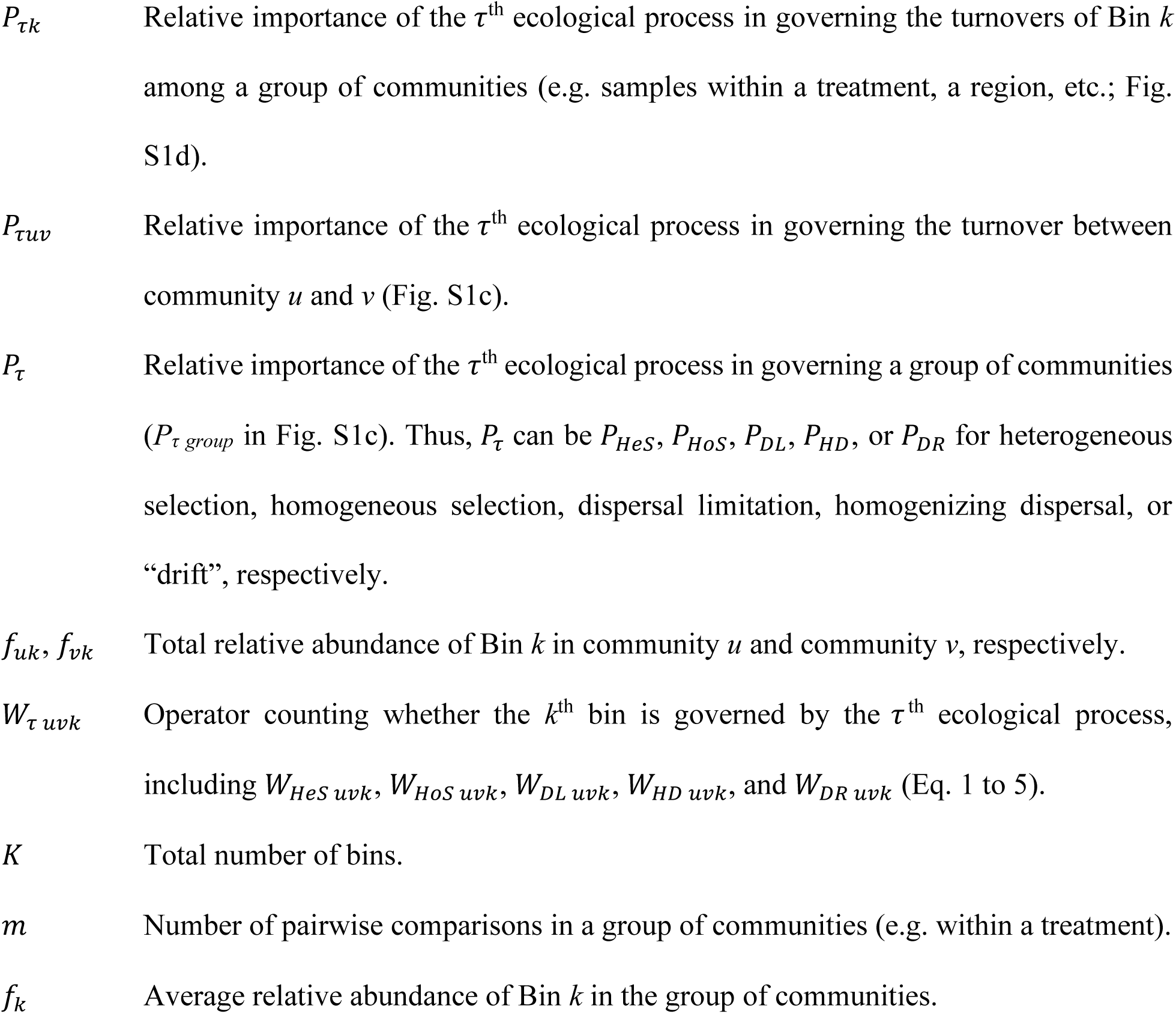

As showed in Eq. 13, the relative importance of each process *P*_*τ*_ is the sum of the terms *f*_*k*_*P*_*τk*_, by which we can define the contribution of different bins to *P*_*τ*_ (Eq. 14 and 15).

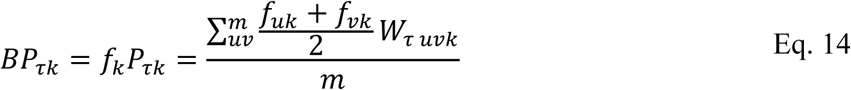

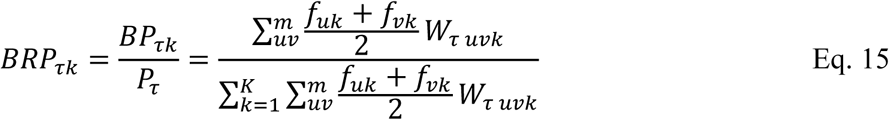

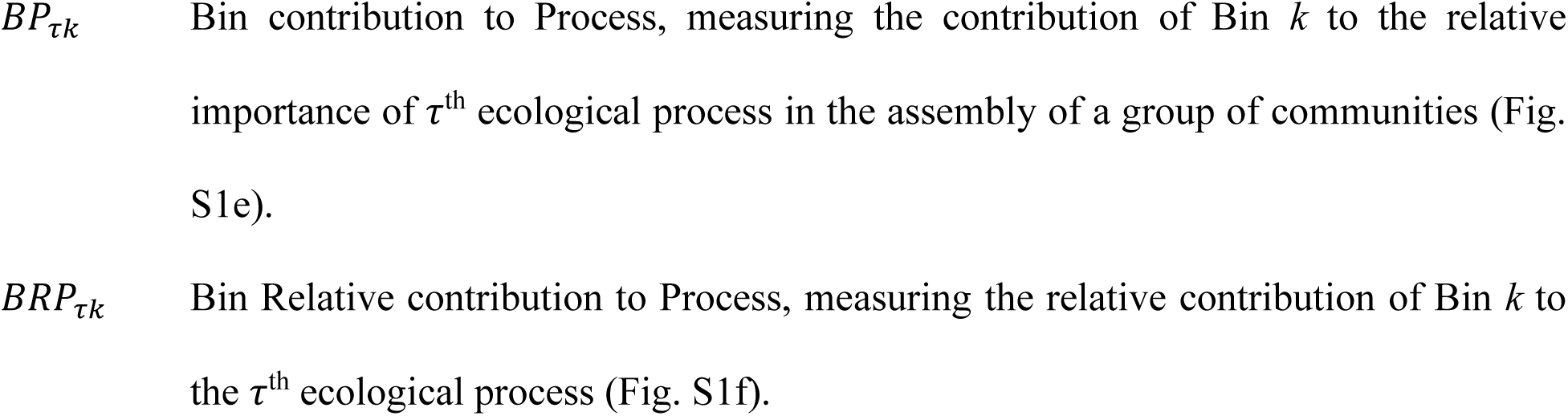

### Simulation model

In the simulation model (Fig. S2), all samples are from the same region sharing the same metacommunity (the regional species pool) with 20 million individuals. The relative abundances of species in metacommunity are simulated using metacommunity zero-sum multinomial distribution model (mZSM) derived from Hubbell’s Unified Neutral Theory Model^60^, using R package “sads”^61^ with J = 2 × 10^7^ and θ = 5,000. The whole region has two separated islands of A and B (Fig. S2a). For species controlled by dispersal, migration is unlimited within each island but nearly impossible between islands. Each island has two plots: plot LA and HA at island A, and plot LB and HB at island B. The two plots at the same island are under distinct environments. The environment variable is as low as 0.05 in the north plots at each island (LA and LB), but as high as 0.95 in the south plots (HA and HB), which is a critical setting for species under niche selection. At each plot, 6 local communities are simulated and sampled as biological replicates. Each local community contains 20,000 individuals of 100 species.

A phylogenetic tree was retrieved from a previous publication^22^ which simulated evolution from a single ancestor to the equilibrium between speciation and extinction and generated a tree with 1,140 species. A trait defining the optimal environment of each species (*E*_*i*_) evolves along the phylogenetic tree with a certain phylogenetic signal. We simulated three pools of species as three scenarios to explore the performance of iCAMP under distinct levels of phylogenetic signals. (i) The low-phylogenetic-signal pool was generated as described previously^22^. The Blomberg’s K value is as low as 0.15, close to the mean K value of 91 continuous prokaryotic traits^44^. The phylogenetic signal is low if counting the phylogenetic distance across the whole tree. However, the trait still shows significant phylogenetic signal within a short phylogenetic distance^22^, in accordance with general observations in microbial communities in various environments^21,40^. (ii) The medium-phylogenetic-signal pool was generated by simulating the trait according to Brownian motion model, using the function “fastBM” in R package “phytools”^62^ with an ancestral state of 0.5, an instantaneous variance of Brownian process of 0.25, and the boundary from 0 to 1. The final K value is 0.9, close to the mean phylogenetic signal level of 899 prokaryotic binary traits^44^. (iii) The high-phylogenetic-signal pool was simulated according to Blomberg’s ACDC model^63^ with a g value of 2000. The final K value is as high as 5.5, close to the highest phylogenetic signal of prokaryotic traits to date^44^.

For each scenario, we simulated 15 situations with different levels of expected relative importance of various processes (Fig. S2b). The situations can be classified into two types. In the first type, all species under each situation are governed by the same kind of processes, i.e. pure selection, or dispersal, or drift. In each of the other situations, species in the regional pool are assigned to different types controlled by various processes. Once a species is assigned to be controlled by selection or dispersal rather than drift, its nearest relatives within *d*_*s*_ will also be assigned to the same type of processes considering the phylogenetic signal of traits. Species controlled by each type of processes are simulated as below. (i) To simulate strong selection without stochasticity, the relative abundance of each species is determined by the difference between the environment variable and their trait values (optimal environment), following a Gaussian function (Eq. 16, Fig. S2d).

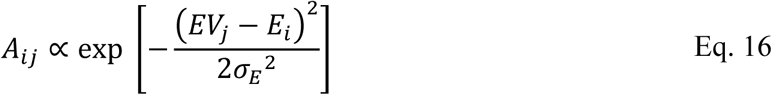

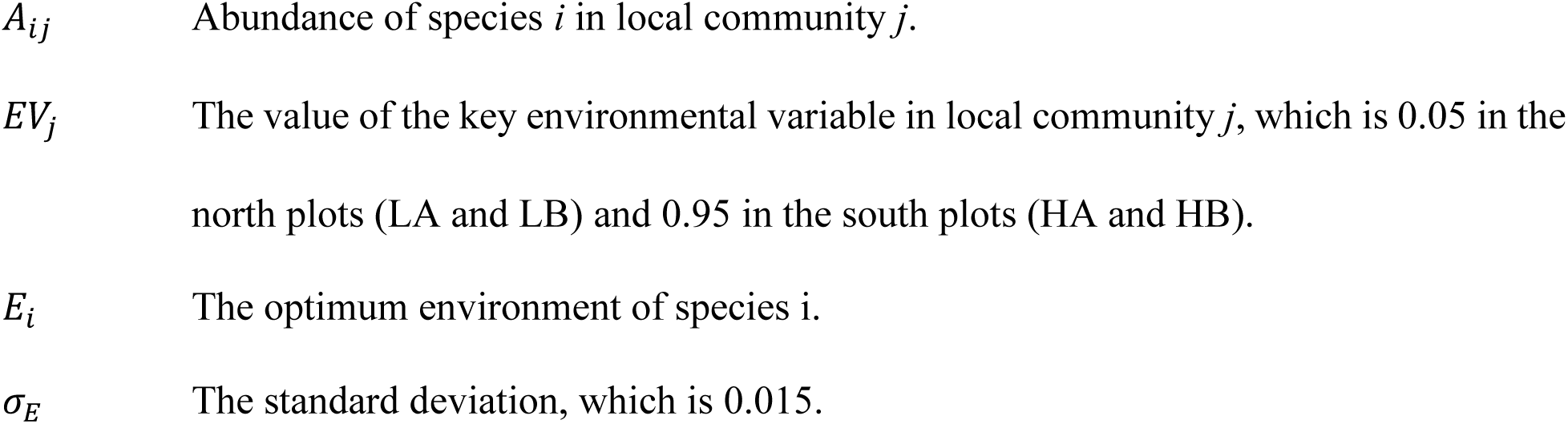

Consequently, the turnovers of these species under the same environment (i.e. within north plots, or within south plots) are solely governed by homogeneous selection, and those between distinct environments (i.e. between north and south plots) are governed by heterogeneous selection.

(ii) To simulate extreme dispersal without selection, we modified Sloan’s simulation model^64^ which was derived from Hubbell’s neutral theory model (Fig. S2e). Each island has a unique species pool, simulated as a local community under the regional metacommunity following neutral theory model but with a relatively low dispersal rate (m_1_ = 0.01). However, the unique species pools of the two islands are constrained to have no overlapped species, regarding extreme dispersal limitation between the two islands. Then, the local communities in each island are simulated as governed by neutral dispersal from both the regional metacommunity with a low rate (m_1_ = 0.01) and the unique species pool of the island with a high rate (m_2_ = 0.99). It means 99% of dead individuals in a local community are replaced by species from the small island-unique species pool at each time step. Therefore, all the turnovers within an island are governed by homogenizing dispersal, and those between islands are controlled by dispersal limitation.

(iii) Drift is simulated as neutral stochastic processes without dispersal limitation or homogenizing dispersal. To simulate drift, all local communities are generated under neutral dispersal from the regional metacommunity with a medium rate (m_1_=0.5, Fig. S2c). Since 50% of dead individuals are replaced by species randomly drawing from a relatively large regional pool, all the turnovers among local communities are neither affected by homogenizing dispersal nor under dispersal limitation.

Under each situation, the dataset of the 24 local communities is simulated as a combination of species governed by different ecological processes, with ratios defined by the situation setting (Table S1, Fig. S2b). For each turnover between a pair of local communities, the mean relative abundance of species governed by a process defines the expected relative importance of the process (Eq. 17). The process with the highest relative importance is the expected dominant process of the turnover. Since dispersal and drift are simulated as pure stochastic processes, the expected stochasticity is defined as the sum of expected relative importance of HD, DL, and DR (Table S1).

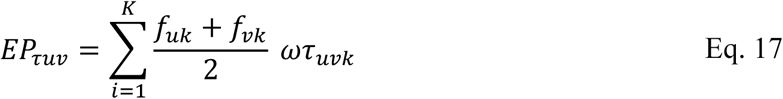

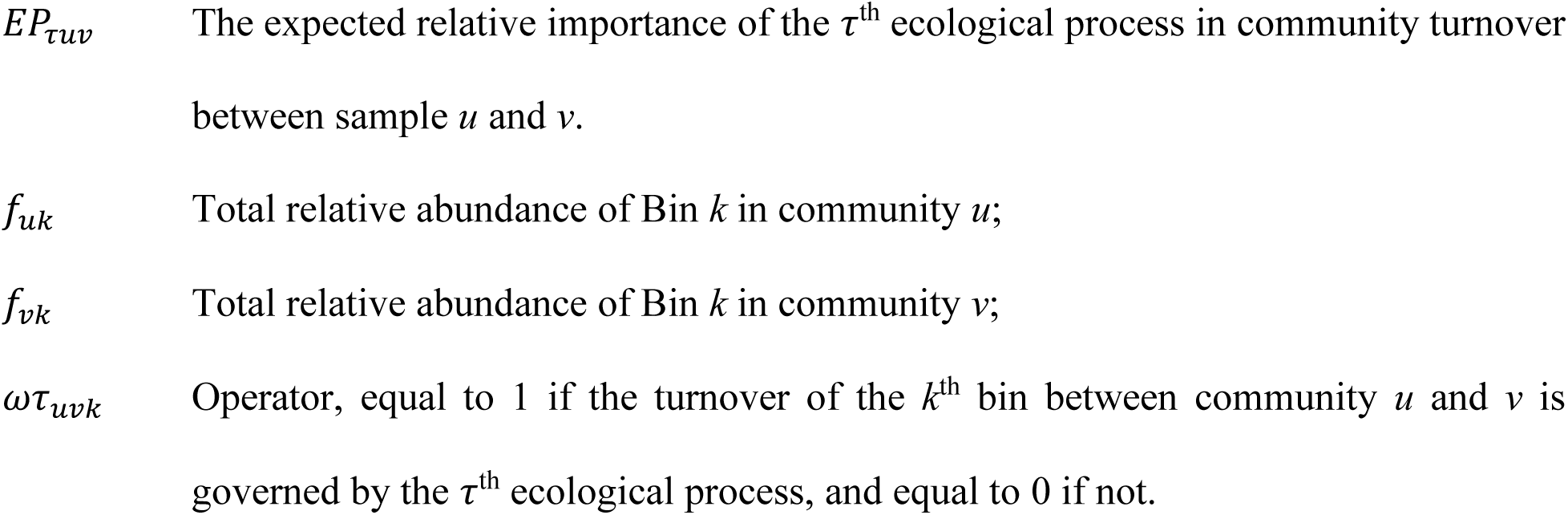

We simulated 3 scenarios with different levels of phylogenetic signal, 15 situations per scenario with 1 dataset per situation, thus a total of 45 datasets. In each dataset, we applied both QPEN and iCAMP to estimate the relative importance of different processes (quantitative estimation) and the dominant process (qualitative estimation). QPEN cannot assess relative importance of processes for each turnover, but can estimate their relative importance as the percentage of turnovers governed by the process in all turnovers within a plot (e.g. plot HA) or between a pair of plots (e.g. plot HA vs HB). Then, the ecological stochasticity of community assembly can be quantified as the relative importance of stochastic processes (i.e. HD, DL, and DR) based on QPEN and iCAMP, respectively. For comparison, the ecological stochasticity in each dataset is also estimated with NP^65^, tNST^37^, and pNST^37,38^, as previously described.

The performance of quantitative estimation is evaluated by accuracy (Eq. 18) and precision coefficients (Eq. 19) derived from Concordance Correlation Coefficient (CCC)^66^. The performance of qualitative estimation is assessed with respect to accuracy (Eq. 20), precision (Eq. 21), sensitivity (Eq. 22), and specificity (Eq. 23) by counting the true and false positive/negative results.

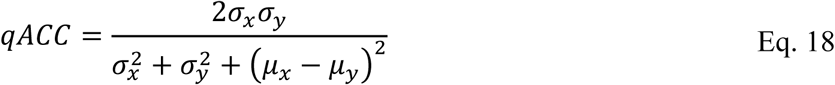

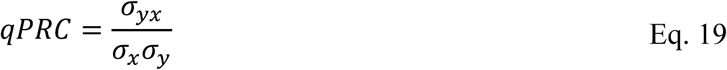

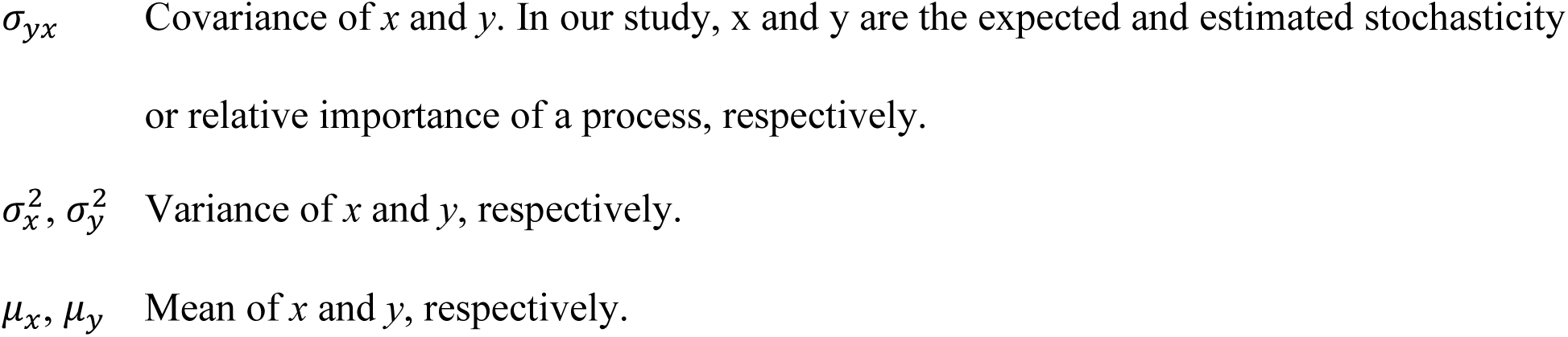

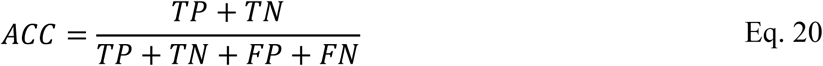

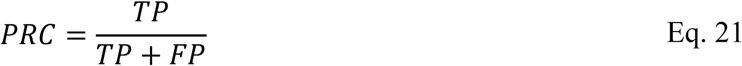

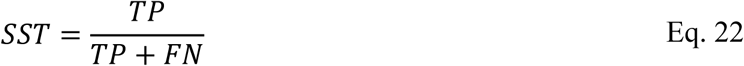

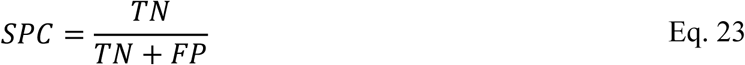

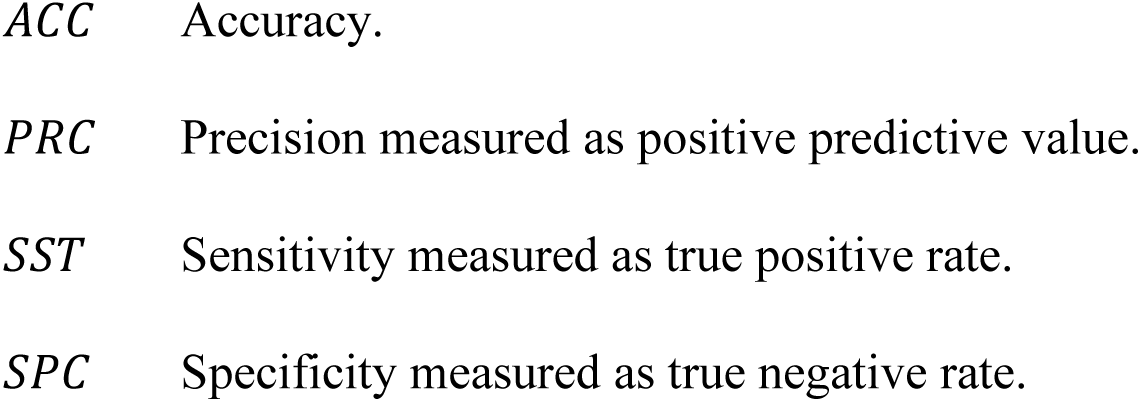
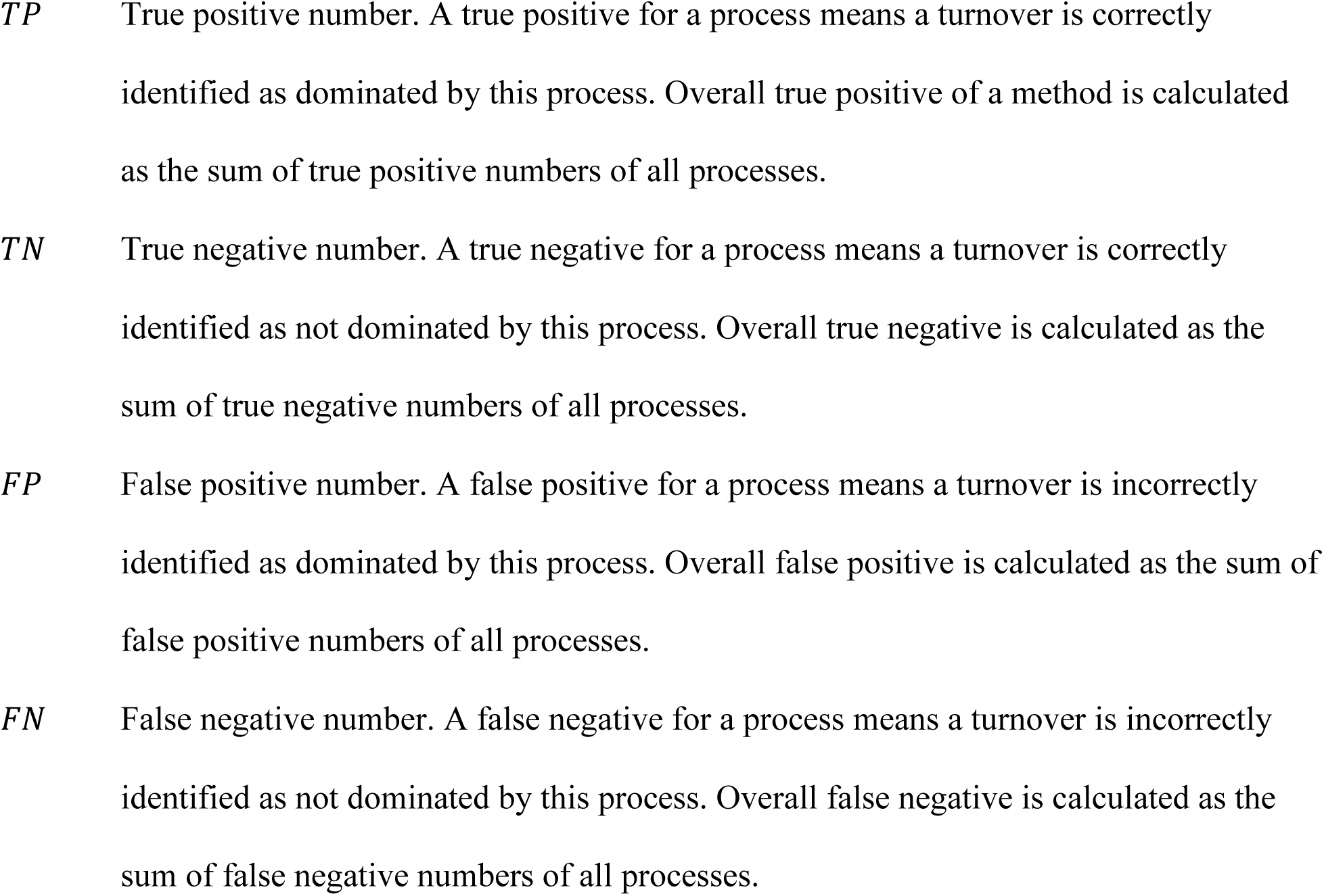

For example, a turnover is in fact dominated by HoS. If the estimated dominating process is HoS, this is a true positive for HoS, and a true negative for other processes. If the estimated dominating process is DL, this is a false positive for DL and a false negative for HoS, but a true negative for HeS, HD, and DR.

### Experimental data and analyses

We applied iCAMP to an empirical dataset from our previous study^38^, with sequencing data available in the NCBI Sequence Read Archive under project no. PRJNA331185. Briefly, the grassland site is located at the Kessler Atmospheric and Ecological Field Station (KAEFS) in the US Great Plains in McClain County, Oklahoma (34° 59′ N, 97° 31′ W)^38^. The field site experiment was established in July of 2009. Surface soil temperature in warming plots (2.5 m × 1.75 m each) is increased to 2 to 3°C higher than the controls by utilizing infrared radiator (Kalglo Electronics, Bath, PA, USA). Surface (0-15 cm) soil samples were taken annually from 4 warming and 4 control plots. A total of 40 samples over 5 years after warming (2010 to 2014) were analyzed in this study. Soil DNA was extracted by from 1.5 g of soil by freeze-grinding and SDS-based lysis as described previously^67^ and purified with a MoBio PowerSoil DNA isolation kit (MoBio Laboratories). DNA was analyzed for bacteria by 16S rRNA gene amplicon sequencing at Illumina MiSeq DNA sequencer, and the sequencing results were processed as described previously^38^. Soil chemistry and plant diversity were measured as previously described^38^. The drought index is calculated as additive inverse of Standardized Precipitation-Evapotranspiration Index (SPEI) retrieved from SPEIbase^68^.

### Statistical analyses

The significance of difference for each evaluation index (e.g. qualitative accuracy, precision, sensitivity, etc.) between different methods was calculated by bootstrapping for 1,000 times (one-side test). To assess the effects of warming on ecological processes, the standardized effect size (Cohen’s d) was calculated as the difference of means between warming and controls divided by the combined standard deviation, and the magnitude of effect is defined as large (|*d*| > 0.8), medium (0.5 < |*d*| < 0.8), small (0.2 < |*d*| < 0.5), and negligible (|*d*| < 0.2) according to Cohen’s *d*^69^. NST^37^, NP^65^ and QPEN^21,22^ were applied to the dataset as previously reported. The significance of difference in stochasticity or relative importance of ecological processes between warming and control was calculated by permutational *t* test (1,000 times). This study only investigated spatial turnovers at each time point.

For correlation test between each process and various measured factors, we applied Mantel test^70^ and MRM^71^ with constrained permutation considering repeated measures design of the experiment. For Mantel test, both linear model and general linear model with a logit link function and a “quasibinomial” distribution were tested, and the relative importance of each process and each factor were either log-transformed or not, to explore the best model. To log-transform a factor with zero or negative values, all its values were subtracted by the lowest value and the resulted zero values were replaced by 0.05 (i.e. −3.00 in natural log) of the minimum positive value before natural -log transformation. For MRM, the factors were forward selected based on adjusted R square. For each measurement (e.g. soil nitrate), both the difference (e.g. |*Nitrate*_*u*_*-Nitrate*_*v*_*|*, where *u* and *v* represent samples) and the mean (e.g. [*Nitrate*_*u*_*+Nitrate*_*v*_]/2) in each pair of samples were investigated for correlation with the relative importance of each process (e.g. *P*_*HoS uv*_). All statistical analyses were implemented by R^73^. All significance tests are two-side unless specified.

## Data Availability

The sequencing data are available in the NCBI Sequence Read Archive under project no. PRJNA331. iCAMP can be implemented using “iCAMP” function on our pipeline (http://ieg3.rccc.ou.edu:8080) and the scripts are available in R package “iCAMP”.

## Acknowledgements

The development of the theoretical framework was supported by the U.S. Department of Energy, Office of Science, Office of Biological and Environmental Research (DOE-BER) Genomic Science program under award number DE-SC0014079 and DE-SC0016247, and also part of ENIGMA-Ecosystems and Networks Integrated with Genes and Molecular Assemblies (http://enigma.lbl.gov), a Scientific Focus Area Program at Lawrence Berkeley National Laboratory, supported by DOE-BER under contract number DE-AC02-05CH11231. The experimental data was generated with the support from DOE-BER under award number DE-SC0010715.

## Author contributions

All authors contributed the intellectual input and assistance to this study and manuscript preparation. J.Z. conceived the research questions. D.N. and J.Z. developed the mathematical framework. M.Y., X.G., and X.Z. collected the empirical data. D.N. developed simulated communities and performed statistical analysis. J.Z. and D.N. wrote the paper with inputs from M.Y., L.W., Y.Z., X.G., Y.Y., A.A., and M.F.

## Competing Interests

The authors declare no competing interests.

