## Supplementary information for "A quantitative framework reveals the ecological drivers of grassland soil microbial community assembly in response to warming"

### Supplementary Text

#### A. Algorithm and parameter optimization for iCAMP

**Phylogenetic binning algorithm.** In the first step of iCAMP, all taxa were divided into bins according to a phylogenetic signal threshold ($d_s$ = 0.2) within which the phylogenetic signal of microbial niche preference was generally found significant in various environments[1-3]. Three phylogenetic binning algorithms were compared (Fig. S3a, b, c), based on phylogenetic distances to abundant species, pairwise phylogenetic distances, and phylogenetic tree, respectively. The algorithms do not have substantial difference in principle, thus led to very similar performances (< 9% difference in performance indexes) of iCAMP when applied to simulated communities (Fig. S1). Under the low- and medium-phylogenetic-signal scenarios, which are more common in the real world, the algorithm based on phylogenetic tree showed slightly higher quantitative precision (up to 3.9% higher, $p < 0.1$) and qualitative performance (up to 8.5% higher) than the other two algorithms (Fig. S3d, e), and thus the tree-based binning is used for iCAMP. This is probably because the relatedness in phylogenetic tree cannot be fully represented by distances. On the contrary, under the high-phylogenetic-signal scenario, the algorithm based on pairwise phylogenetic distance showed slightly better performance than other algorithms (Fig. S1f). Considering microbial traits in the real world usually have low or medium phylogenetic signal[4], the tree-based algorithm is more recommended for iCAMP.

**Minimal bin size.** To maintain statistical power within each bin, a minimal requirement of bin size (minimal taxa number in a bin, $n_{min}$) should be defined. If a bin has too few taxa, it will be merged into the most relevant bin. However, high $n_{min}$ may make the phylogenetic distances within some bins too large to maintain phylogenetic signal. A series of the values of $n_{min}$ from 6 to 96 were compared for their impacts on iCAMP performance. While quantitative accuracy was not significantly affected, all other indexes, especially quantitative precision, qualitative precision, and sensitivity of iCAMP, were significantly ($p < 0.05$) influenced by $n_{min}$, with the best performance at $n_{min}$ = 24 (Fig. S4a, b, c). To explore the reason, the phylogenetic signal within each bin was analyzed by Mantel tests between phylogenetic distance and niche difference under different $n_{min}$. When $n_{min}$ = 24, the bins with significant (one-tail $p < 0.05$ and R > 0.10, since R < 0.10 is usually regarded as negligible effect size[5]) phylogenetic signal reached the highest relative abundance (Fig. S4d, e, f). When other $n_{min}$ values showed the same relative abundance of significant bins (e.g. $n_{min}$ = 48 and 96 in Fig. S4e, f), $n_{min}$ = 24 led to higher effect size, measured as average R value of within-bin phylogenetic signal. Thus, $n_{min}$ could be determined according to within-bin phylogenetic signal, i.e. higher relative abundance of bins with significant phylogenetic and higher R value of within-bin phylogenetic signal.

**Phylogenetic metrics.** To identify the impacts of selection, the phylogenetic metric βNTI (beta Nearest Taxon Index) has been widely used in recent studies[2,3,6-10], mainly because βNTI is based on βMNTD (beta Mean Nearest Taxon Distance) of which the values are generally within the phylogenetic signal threshold[7,10], and thus it is better than other phylogenetic metrics to reflect niche preference dissimilarity. However, in iCAMP, the phylogenetic distances in each bin are mostly within the phylogenetic signal threshold, thus, another phylogenetic metric βNRI (beta Net Relatedness Index) based on βMPD (beta Mean Pairwise Distance) is also applicable. When applied to the simulated communities, βNRI resulted in obviously ($p < 0.01$) higher quantitative precision (13.8-17.3% higher) and qualitative performance (2.7-28.6% higher) of iCAMP than βNTI did under medium- and high-phylogenetic-signal scenarios (Fig. S5b, c), but only slightly higher quantitative precision (3% higher) than βNTI under low-phylogenetic scenario (Fig. S5a). While βNTI only counts the distance of each taxon to its nearest relative, βNRI counts distance of each taxon to all other taxa, and hence it includes more information than βNTI. The higher the across-tree phylogenetic signal is, the more useful information βNRI can count in than βNTI. Therefore, the advantage of βNRI in iCAMP is more obvious under scenarios with higher phylogenetic signal. Accordingly, βNRI is preferred in iCAMP.

**Randomization range in null models.** Besides phylogenetic metrics, the null model algorithm is also critical in the second step of iCAMP. As in QPEN, the phylogenetic null model is utilized to infer selection, and taxonomic null model is used to further identify dispersal limitation and homogenizing dispersal[3,11]. For both null models, the randomization can be performed within each bin or across taxa in all bins. In the phylogenetic null model for βNRI calculation, within-bin randomization led to higher quantitative precision and qualitative performance (1.6-16.4% higher, $p < 0.05$) of iCAMP than across-bin randomization, especially under low-phylogenetic-signal scenario (Fig. S6a, b, c). This is mainly due to significant phylogenetic signal within bin rather than across bins, which is important for using βNRI to infer selection. In contrast, the taxonomic null model using across-bin randomization resulted in obviously higher quantitative precision and qualitative performance (8.6~103% higher, $p < 0.05$) for iCAMP than using within-bin randomization (Fig. S6d, e, f). This is reasonable considering that the taxonomic null model analysis is used to infer neutral dispersal process, which is not species-specific but influences taxa across all bins in probability as long as under the same metacommunity. Therefore, for iCAMP, βNRI should be calculated based on within-bin randomization, and RC should be estimated based on across-bin randomization.

**Randomization times.** The null model analysis needs enough randomization times to estimate the distribution of the null values of phylogenetic/taxonomic dissimilarity. Low randomization times cannot provide reproducible results, but high randomization times cost more computational resources and/or time. Randomization times ranging from 25 to 5,000 were used for iCAMP analysis of simulated communities, and the quantitative and qualitative results of iCAMP were compared with the iCAMP result from 60,000-time randomization. When the randomization times are not less than 200, deviation of quantitative results and the error rate of qualitative results were less than 0.05 (down to zero) and became relative stable (no significant change in mean and deviation) as randomization times increased (Fig. S7). Thus, the commonly used randomization times (1,000 times) are generally enough for iCAMP analysis, and 200 times may also be acceptable for relatively small data (e.g. less than 2,000 OTUs).

**Reducing the taxa number.** Besides randomization times, a large taxa number can cause quadratic increase of computational resource demand and time cost for phylogenetic null model analysis, making iCAMP and QPEN much less feasible for very large dataset (e.g. over 100,000 OTUs). In addition, the taxa with low relative abundances may bring more technical noise, for example, in the OTU tables from amplicon sequencing[12-14]. Thus, large data may need to be reduced before the iCAMP analysis. Three methods were compared, including classic rarefaction (rarefaction), cutting based on average relative abundance across samples (average abundance cut), and cutting based on cumulative abundance in each sample (cumulative abundance cut). Their performance was evaluated according to the quantitative deviation and qualitative error rate of iCAMP using the reduced OTU table compared to results from original OTU table. Rarefaction led to obvious overestimation of the DR importance (Fig. S8), and the error rate of qualitative estimation can be larger than 90% even though over 70% species were still remained (e.g. qualitative in Fig. S8a). This is probably because rarefaction itself is a random sub-sampling process bringing more artificial stochasticity, and also removes much more sequences than other methods to reduce taxa number. The other two methods resulted in similar performance of iCAMP (Fig. S8). After cumulative abundance cut, quantitative deviation was usually lower than 10% when taxa number was reduced to a half of the original number, but can be up to 87% (e.g. for HoS in Fig. S8b) at 30% of the original taxa number. After abundance cut, qualitative error rates were generally more than 10%, in some cases up to 60%. Altogether, to investigate assembly mechanism, sequencing should be deep enough with high coverage to minimize the negative impact of random sampling. Reducing taxa number can significantly change the estimation, and thus it is not advisable, except necessary denoising, i.e. removing unreliable technical noise (e.g. removing global singleton)[12]. Our program applies parallel computation, matrix-based algorithm, and “big memory”[15] (efficiently utilizing hard disk as memory) to make iCAMP feasible for a relative large dataset (10,000-40,000 OTUs) on a desktop computer or a small server. For a very large dataset, high performance computing can help, if not feasible, the taxa number can be reduced by cumulative abundance cut before iCAMP analysis. Although the quantitative results could be acceptable, the qualitative results are not reliable.

**B. Linkages between environmental factors and ecological processes**

The relative importance of the two major processes (HoS and DR) was analyzed for their linkages to all measured environmental factors and spatial variables by Mantel test and Multiple Regression on distance Matrices (MRM). Since the relative importance of other processes (HeS, HD, and DL) appear small, we will focus on HoS and DR.

In control plots, the HoS importance showed significant (Mantel, Fig. 5a) negative correlation with precipitation (level of precipitation in sampling month, $p < 0.01$), soil moisture (variation of annual mean soil moisture, $p < 0.01$), and plant richness (level of plant richness, $p < 0.01$), but positive correlation with drought (level of the drought index in sampling month, $p < 0.05$) and C4 plant biomass (variation of aboveground biomass of C4 plant, $p < 0.01$), with similar strength ($R^2$ = 0.30-

0.34), which were higher than all other factors ($R^2 < 0.16$, Fig. 5a, Table S2). The DR importance also showed significant correlation with these factors, but in an opposite way, i.e. positively with precipitation, moisture, plant richness, but negatively with drought and C4 plant biomass (Fig. 5b). The C4 plant biomass was related to plant diversity, since plants were usually less diverse when a few C4 plant species grew well and predominated the plant community. These results suggested lower water and plant diversity increased selection and decreased drift, leading to more clustered species in soil bacterial communities.

In MRM of HoS and DR in control plots, plant richness was ranked first (explained 33% variance) for HoS (Fig. S15a), and C4 plant biomass was scored the first (35%) for DR (Fig. S15c). When including other factors in the MRM models of HoS and DR, each factor explained much lower proportions of the variations (explained 0.02–12.2%) than the first one, but cumulatively, other factors added over 65% explanation power to the total $R^2$ very close to 1 (Fig. S15a, c).

In warming plots, drought index in sampling month showed significant, negative correlation with HoS (Fig. 5a) and positive with DR (Fig. 5b), while precipitation showed opposite correlation, with $R^2$ from 0.52 to 0.57, much stronger than those under control, suggesting warming enhanced the filtering effect of drought resulting from low precipitation. Moisture did not show any significant correlation with HoS and DR in warming plots ($R^2 < 0.15$, $p > 0.4$, Table S2), implicating the drought-induced filtering effect in warming plots was different from the moisture-related filtering in control plots. In addition, plant biomass (level of aboveground biomass of C4 and all plants) also showed strong association with HoS (negatively, Fig. 5a) and DR (positively, Fig. 5b) in warming plots, but plant richness did not, suggesting the plant-oriented selection pressure was shifted from lower plant diversity to lower productivity by warming. Soil temperature, which was not correlated with HoS in control plots ($R^2 < 0.031$), shifted to significantly correlated with HoS and DR, however the association was considerably weaker than those with drought and plant biomass. Warming also altered the correlation between soil chemical variables and HoS/DR, e.g. nitrate was correlated with HoS ($p < 0.1$) in controls but not in warming plots, and ammonium was correlated with HoS and DR in warming plots but not in controls. However, all soil chemical variables showed much lower $R^2$ than drought and plant. In multiple regression of HoS and DR in warming plots, precipitation, plant biomass, and soil temperature were the major factors, which explained ~80% of the variance of HoS (Fig. S15b) and DR (Fig. S15d). Altogether, the factors related to water, plant biomass and temperature showed much stronger association with HoS under warming than control conditions, suggesting that climate warming significantly enhanced selection pressure imposed by drought (lower precipitation), lower plant productivity, and increased soil temperature on grassland bacteria, leading to community structure with less variability and stochasticity.

#### Supplementary Figures

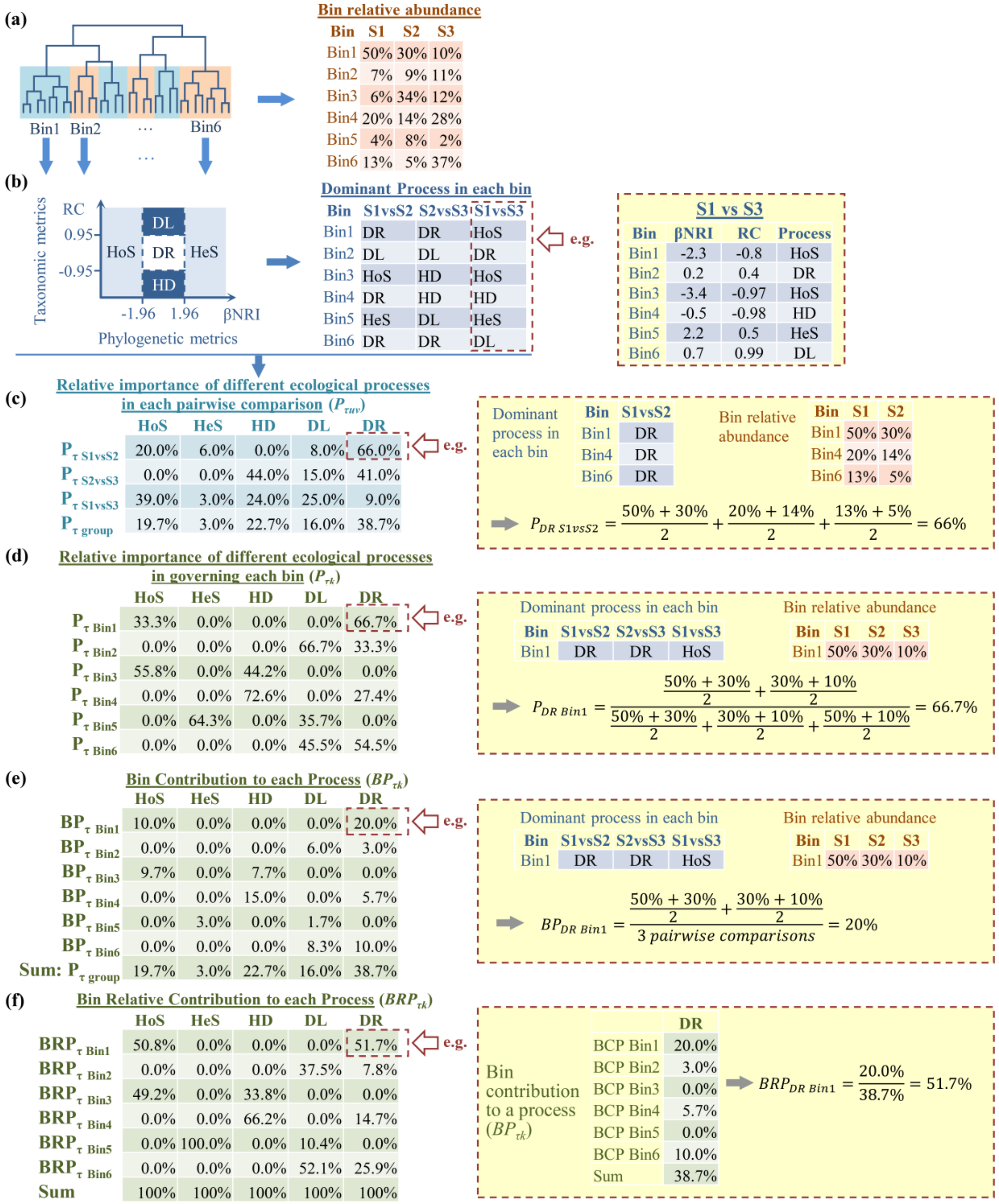


**Fig. S1. A simple example of iCAMP analysis. (a)** Phylogenetic binning based on phylogenetic tree. **(b)** Identifying an ecological process governing the turnover of each bin (Eq. 1-10). **(c)** Relative importance of different ecological processes in governing each pairwise community turnover (Eq. 12, 13). A community turnover means the dissimilarity/similarity found in a pairwise comparison between two communities or samples. **(d)** Relative importance of different ecological processes in governing each bin (Eq. 11). **(e)** Contribution of each bin to each process (Eq. 14). **(f)** Relative contribution of each bin to each process (Eq. 15). Boxes with red dashed lines showed detailed calculation examples. Bin1 to Bin6, phylogenetic bin IDs. S1, S2, S3, sample IDs. HoS, homogeneous selection; HeS, heterogeneous selection; HD, homogenizing dispersal; DL, dispersal limitation; DR, "drift" including stochastic drift, diversification, weak selection and/or weak dispersal. βNRI, beta net relatedness index calculated from beta mean pairwise distance (βMPD); RC, modified Raup-Crick metrics based on Bray-Curtis dissimilarity.

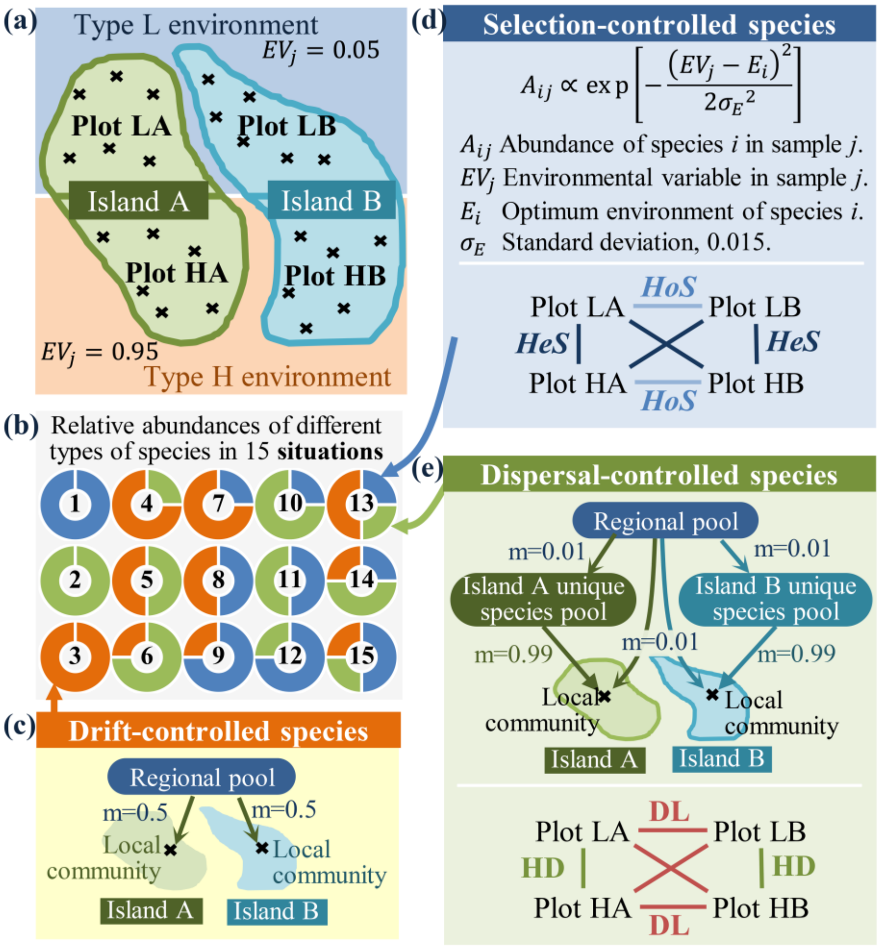


**Fig. S2. Illustration of the basic settings of the simulated communities. (a)** Predefined locations of sampling plots (LA, LB, HA, HB) and local communities (black cross) in two islands (A, B) and two types of environments (L, e.g., low temperature; H, e.g., high temperature). **(b)** Predefined relative abundances of species controlled by various processes under 15 different situations. Blue, selection-controlled species; green, dispersal-controlled species; orange, drift-controlled species. These 15 situations were simulated under each of the three scenarios, i.e. low, medium, and high phylogenetic signal of the key traits. **(c)** Immigration setting to simulate species controlled by drift without too high or low dispersal rate. *m* is dispersal rate in neutral theory model, which is the probability that a dead individual will be replaced by an individual immigrating from the regional pool rather than by a local individual. Since the dispersal (m = 0.5) is neither limited (m → 0) nor homogenizing (m → 1), the turnovers of these species are all controlled by drift. **(d)** Relative abundance of selection-controlled species determined by the key trait ($E_i$). For selection-controlled species, the turnovers between plots under the same environments (Plot LA vs LB and plot HA vs HB) is governed by homogeneous selection (HoS), and those between different environments (LA vs HA, LA vs HB, LB vs HA, and LB vs HB) is governed by heterogeneous selection (HeS). **(e)** Immigration setting to simulate species controlled by dispersal and the dominant processes between different plots. For dispersal-controlled species, the turnover within each island (LA vs HA, LB vs HB) is controlled by homogenizing dispersal (HD), and their turnover between different islands (LA vs LB, LA vs HB, HA vs HB, and HA vs LB) is controlled by dispersal limitation.

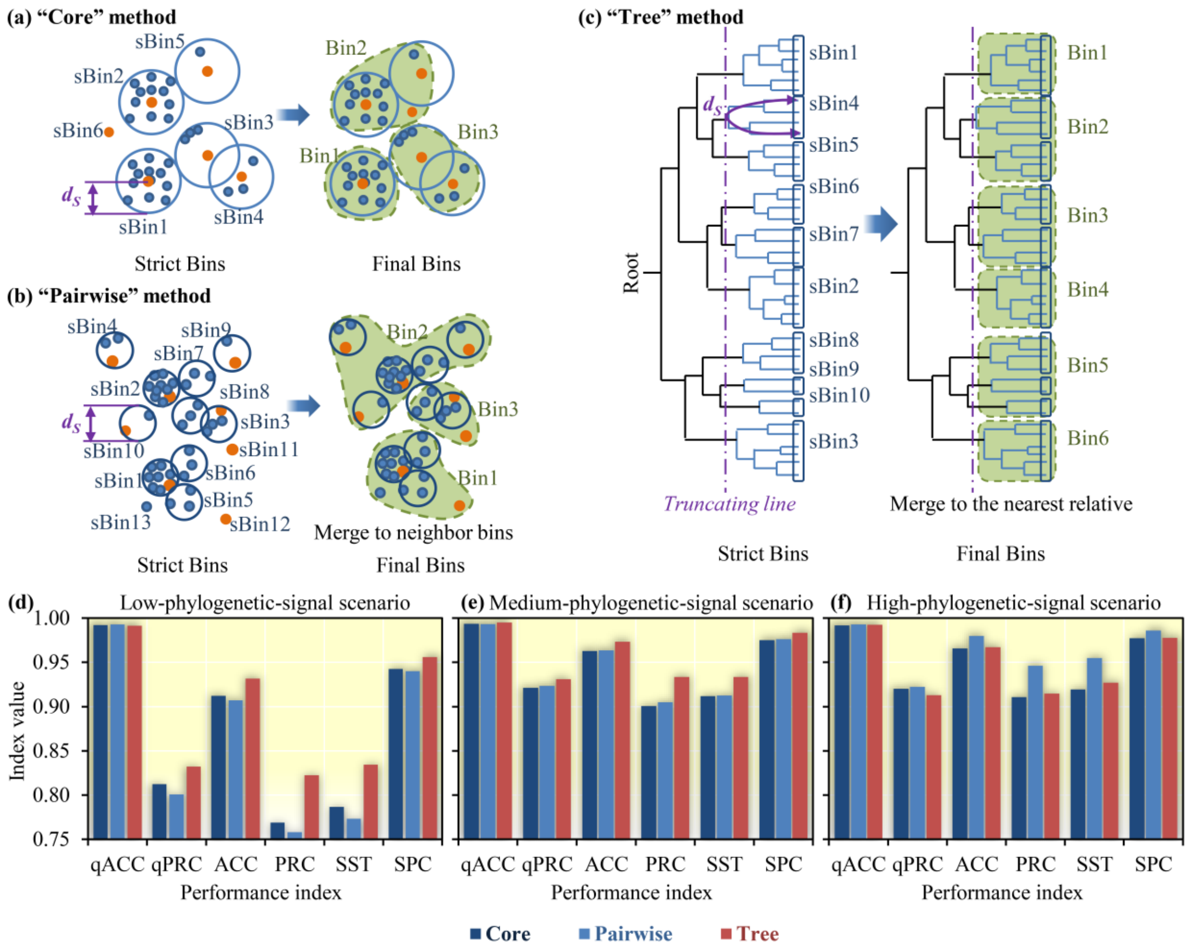


**Fig. S3. Comparison of three methods for phylogenetic binning in iCAMP. (a)** Binning based on phylogenetic distance to abundant species ("**core**" method). In a strict bin (sBin1-6), all distances to the core species (orange dot, the most abundant species) are shorter than the phylogenetic signal threshold ($d_s$). To ensure enough statistical power, bins with too few species are combined with the nearest neighbor(s) until all final bins (Bin1-3) reach the minimal requirement ($n_{min}$, set as 6 in this figure). **(b)** Binning based on pairwise phylogenetic distance ("**pairwise**" method). In a strict bin (sBin1-13), all pairwise distances are shorter than $d_s$. Then, small bins are combined with the nearest neighbor(s) to form final bins with species no less than $n_{min}$. **(c)** Binning based on phylogenetic tree ("**tree**" method). In a strict bin, all species have the same ancestor after the truncating point and all pairwise phylogenetic distances are shorter than $d_s$. Then, small bins are merged to the nearest relative(s) to form final bins. **(d)** Performance of iCAMP using different phylogenetic binning methods under low phylogenetic signal. **(e)** medium phylogenetic signal, and **(f)** high phylogenetic signal. Performance indexes include quantitative accuracy (qACC) and precision (qPRC), qualitative accuracy (ACC), precision (PRC), sensitivity (SST), and specificity (SPC).

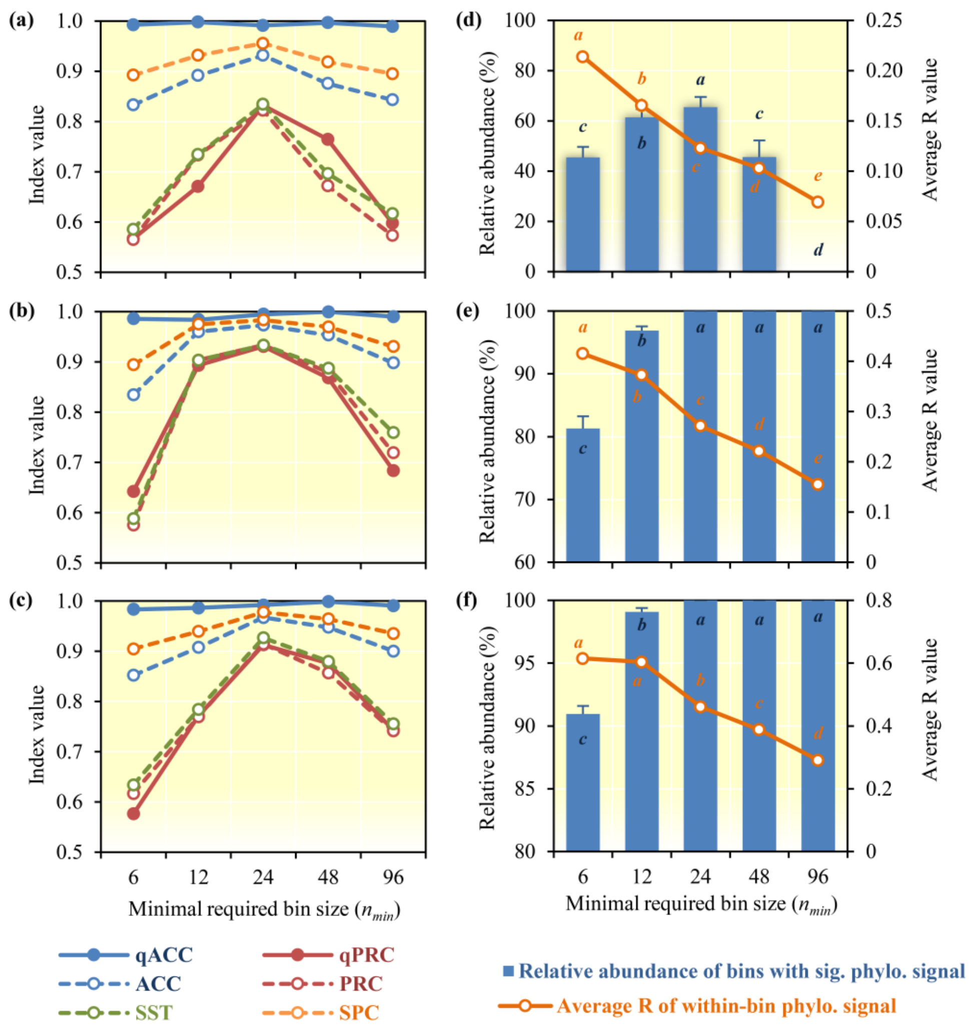


**Fig. S4. Effects of minimum required bin sizes ($n_{min}$) on iCAMP performance and within-bin phylogenetic signal. (a)** Performance of iCAMP using different $n_{min}$ under low-phylogenetic-signal scenario, **(b)** medium-phylogenetic-signal scenario, and **(c)** high-phylogenetic-signal scenario. **(d)** Evaluation of within-bin phylogenetic signal when different $n_{min}$ under low-phylogenetic-signal scenario is used, **(e)** medium-phylogenetic-signal scenario, and **(f)** high-phylogenetic-signal scenario. Here, within-bin phylogenetic signal means correlation between the pairwise phylogenetic distances and niche preference differences among species within the same bin. Within-bin phylogenetic signal is evaluated by relative abundance of bins with significant phylogenetic signal and average R value of all within-bin Mantel test. Performance indices are abbreviated as Fig. 3. In a bin, the phylogenetic signal is regarded as significant if R > 0.10 (R < 0.10 is usually regarded as negligible effect size[5]) and one-tail $p < 0.05$ in Mantel test between phylogenetic distances and niche preference difference (key trait difference). In panel d-f, error bars indicate standard error of the relative abundances under different situations in each scenario, and different letters represent significant difference ($p < 0.05$).

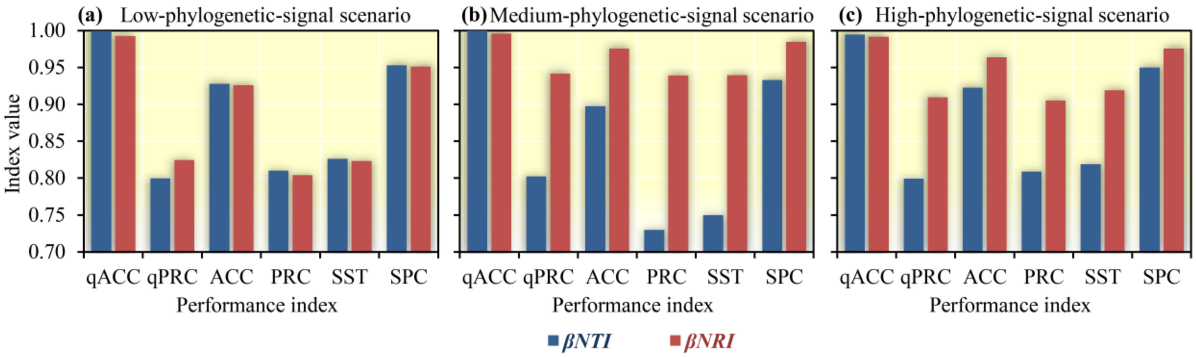


**Fig. S5. Effects of phylogenetic metrics on iCAMP performances under different simulation scenarios.** **(a)** Low-phylogenetic-signal scenario; **(b)** Medium-phylogenetic-signal scenario; **(c)** High-phylogenetic-signal scenario. βNTI, blue; βNRI, red. Performance indexes are abbreviated as Fig. S3.

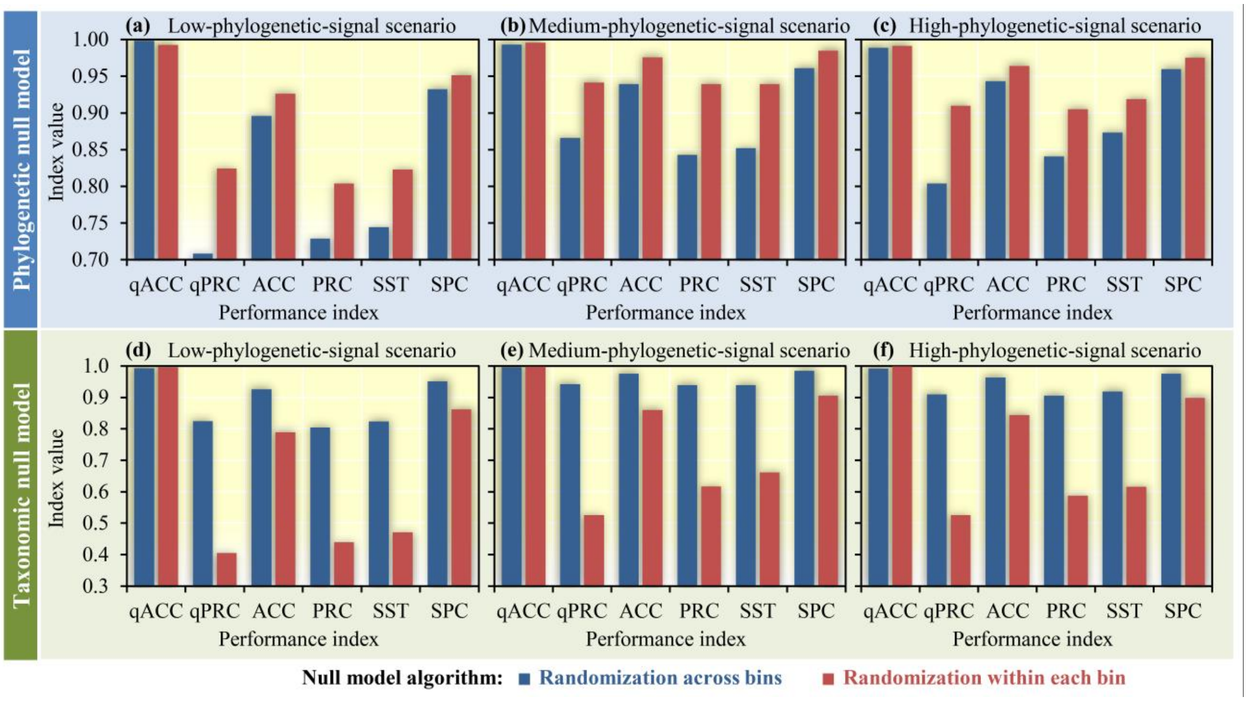


**Fig. S6. Effects of randomization range in phylogenetic and taxonomic null models on iCAMP performances. (a)** Performances of iCAMP using phylogenetic null model with different randomization ranges under low-phylogenetic-signal scenario; **(b)** Medium-phylogenetic-signal scenario; **(c)** High-phylogenetic-signal scenario. **(d)** Performances of iCAMP using taxonomic null model with different randomization ranges under low-phylogenetic-signal scenario; **(e)** Medium-phylogenetic-signal scenario; **(f)** High-phylogenetic-signal scenario. Blue, randomization across bins; red, randomization within each bin. Performance indices are abbreviated as Fig. S3.

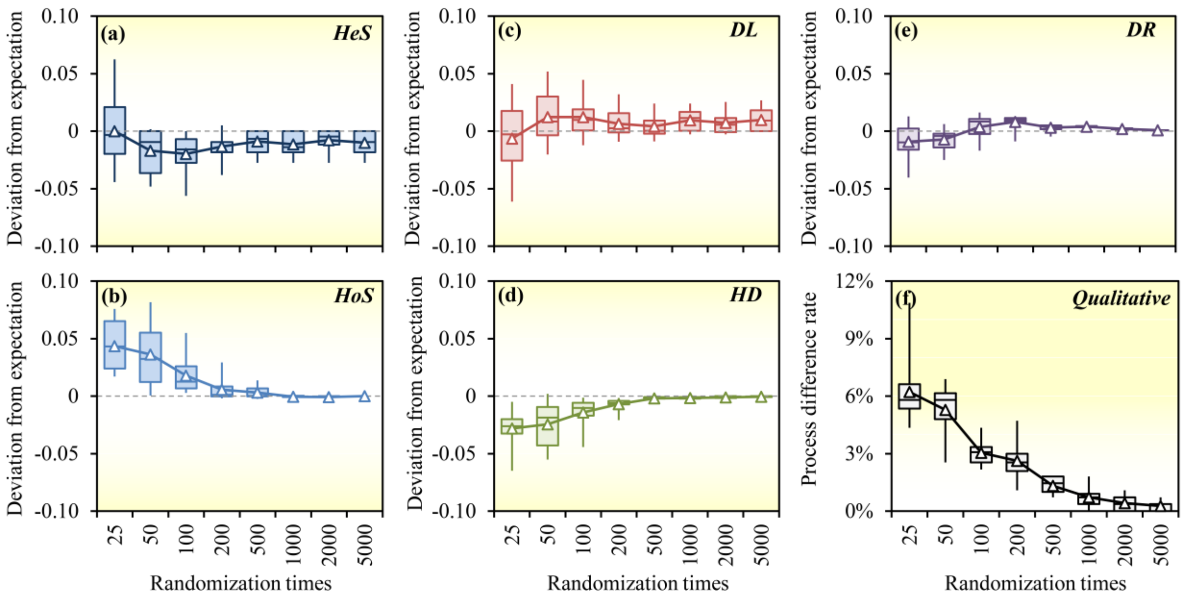


**Fig. S7. Effects of randomization times on the reproducibility of quantitative and qualitative results from iCAMP. (a)** Deviation of the estimated relative importance of HeS from expectation under different randomization times. **(b)** Deviation of HoS. **(c)** Deviation of DL. **(d)** Deviation of HD. **(e)** Deviation of DR. **(f)** The rate of dominant processes estimated differently from expectation under different randomization times. iCAMP was applied to simulated communities under the 12$^{th}$ situation (75% selection, 25% dispersal, Table S1) in low-phylogenetic-signal scenario. Deviation from expectation was calculated as the difference of the estimated relative importance of a process from the result after 60,000-time randomizations. Process difference rate, percentage of the estimated dominated processes which are different from the estimation after 60,000-time randomizations. Box and whisker, quartiles of results from iCAMP repeated for 12 times; triangle, mean value.

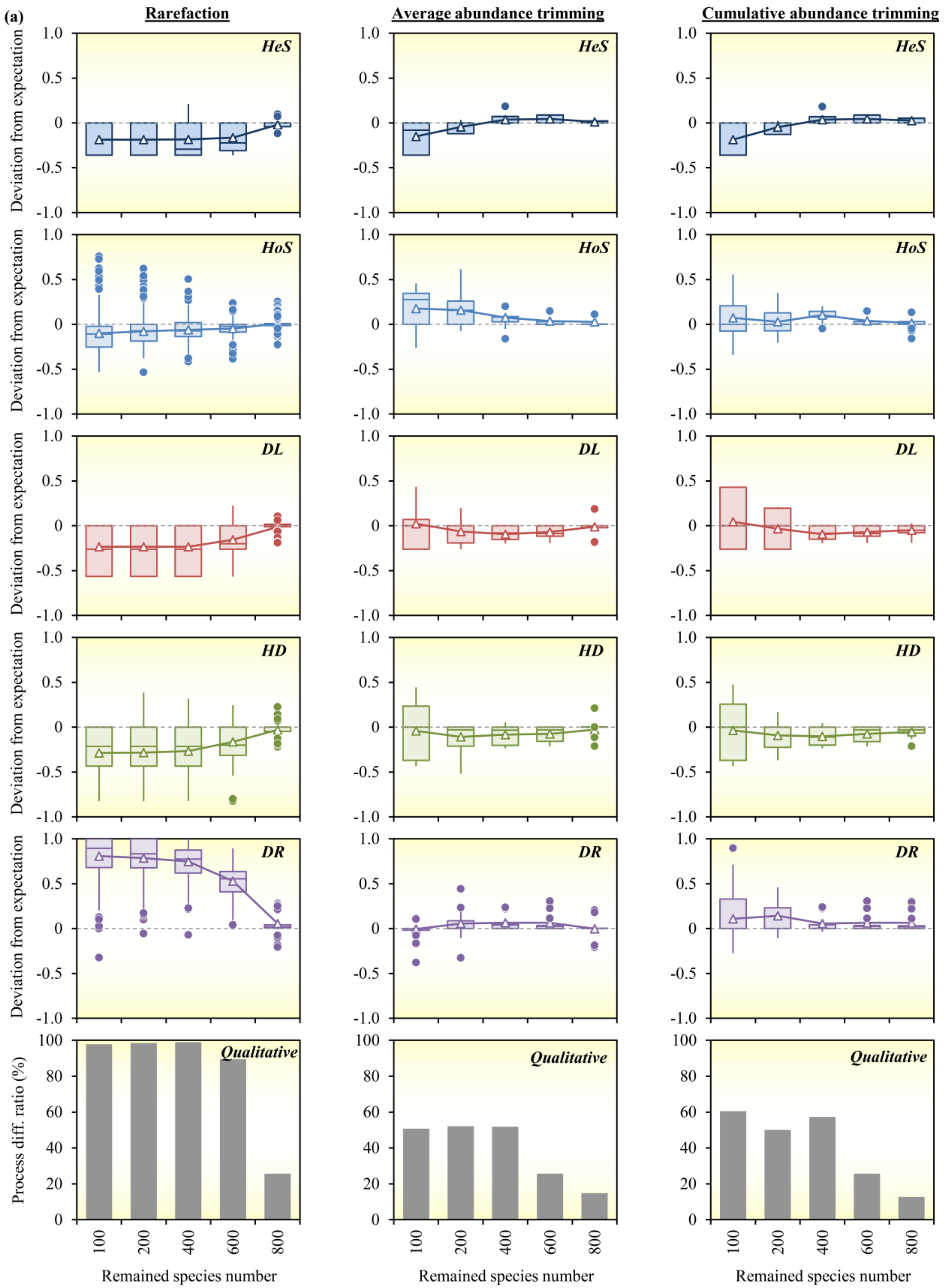
(a)
Rarefaction
Average abundance trimming
Cumulative abundance trimming
HeS
HoS
DL
HD
DR
Qualitative
Deviation from expectation
Process diff. ratio (%)
Remained species number
1.0
0.5
0
-0.5
-1.0
100
80
60
40
20
0
100
200
400
600
800

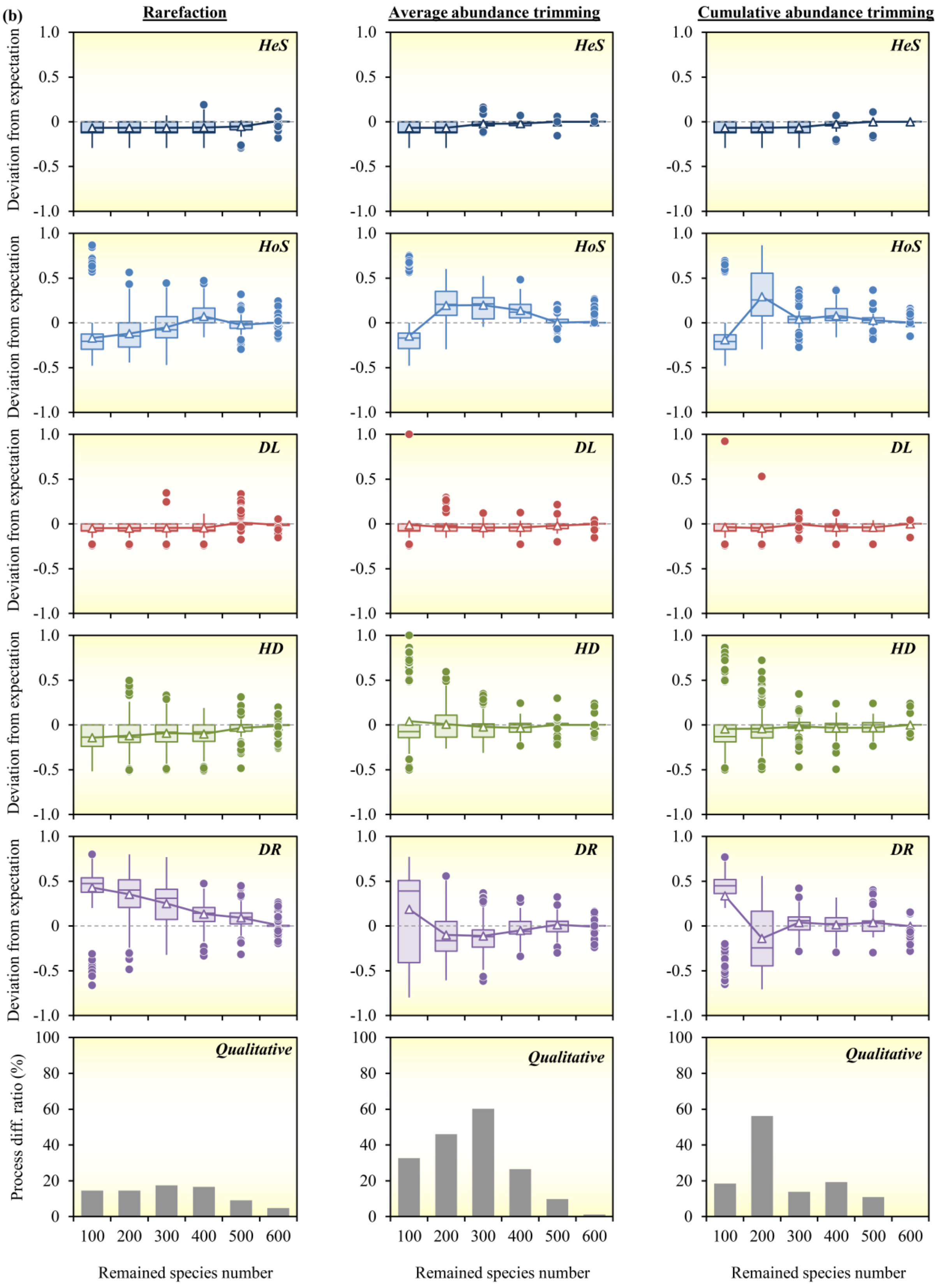

(b)
Rarefaction
Average abundance trimming
Cumulative abundance trimming
HeS
HoS
DL
HD
DR
Qualitative
Deviation from expectation
Process diff. ratio (%)
1.0
0.5
0
-0.5
-1.0
100
80
60
40
20
0
100 200 300 400 500 600
Remained species number

**Fig. S8. Effects of taxa number reduction with different methods on the reproducibility of quantitative and qualitative results from iCAMP. (a)** Results from simulated communities under the 12th situation (75% selection, 25% dispersal, Table S1) in low-phylogenetic-signal scenario. **(b)** Results from simulated communities under the 13th situation (25% selection, 25% dispersal, 50% drift, Table S1) in low-phylogenetic-signal scenario. Deviation from expectation is calculated as the difference of the estimated relative importance of a process from the result of the original data. Process difference rate, percentage of the estimated dominated processes which are different from the results of original data. Box and whisker, quartiles; triangle, mean value.

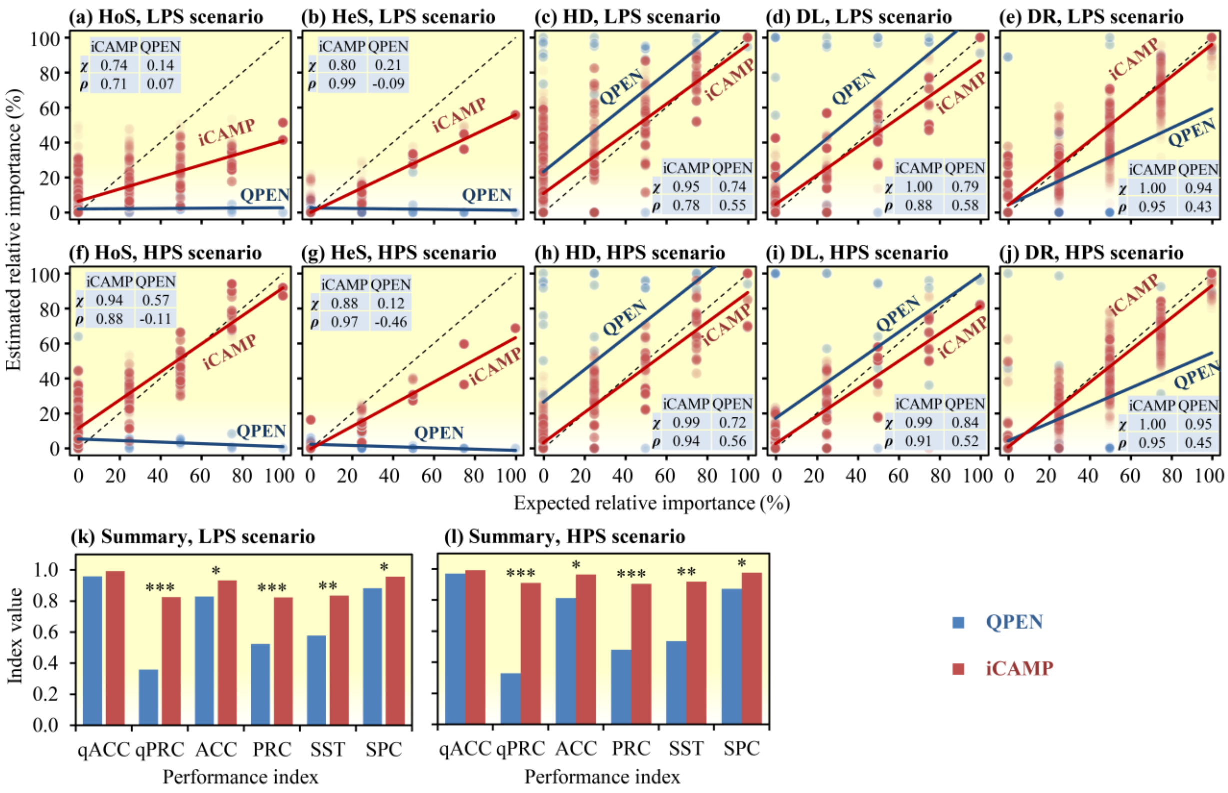


**Fig. S9. Performances of iCAMP (red) and QPEN (blue) with simulated communities.** The performance was evaluated by the consistency of estimated and expected relative importance of individual processes under different phylogenetic signal scenarios: **(a)** HoS under low-phylogenetic-signal scenario (LPS), **(b)** HeS under LPS, **(c)** HD under LPS, **(d)** DL under LPS, **(e)** DR under LPS, **(f)** HoS under high-phylogenetic-signal scenario (HPS), **(g)** HeS under HPS, **(h)** HD under HPS, **(i)** DL under HPS, and **(j)** DR under HPS. **(k)** Overall performance of iCAMP and QPEN under LPS evaluated by six performance indexes: quantitative accuracy (qACC, $\chi$) and precision (qPRC, $\rho$), qualitative accuracy (ACC), precision (PRC), sensitivity (SST), and specificity (SPC); (l) Performance of iCAMP and qPEN under high-phylogenetic-signal scenario (HPS).

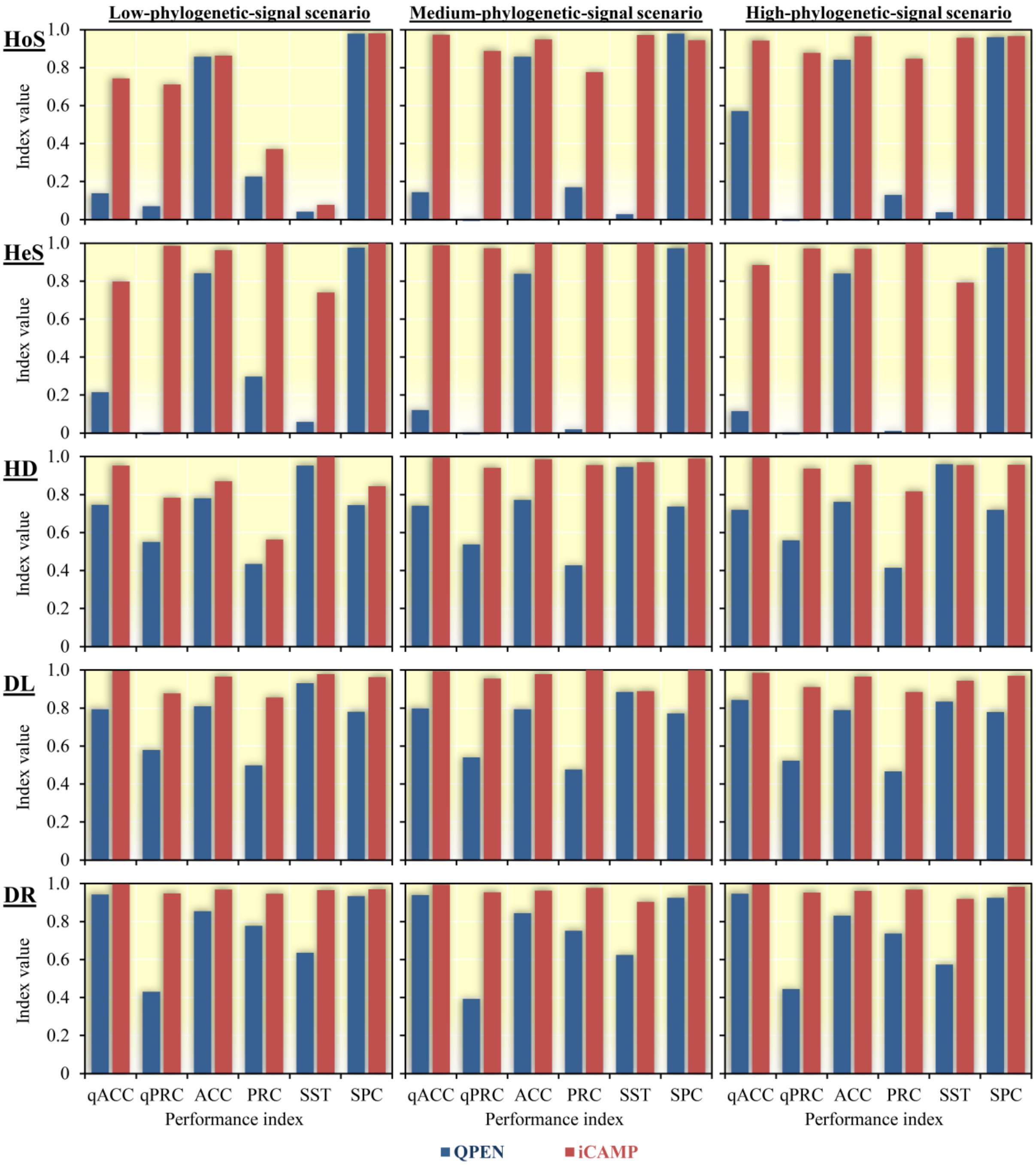


**Fig. S10. Performances of iCAMP (red) and QPEN (blue) in assessing the relative importance of different ecological processes in simulated communities.** iCAMP and QPEN were applied to simulated communities under different scenarios with low, medium, and high phylogenetic signals. Performance indices were abbreviated as in Fig. S3.

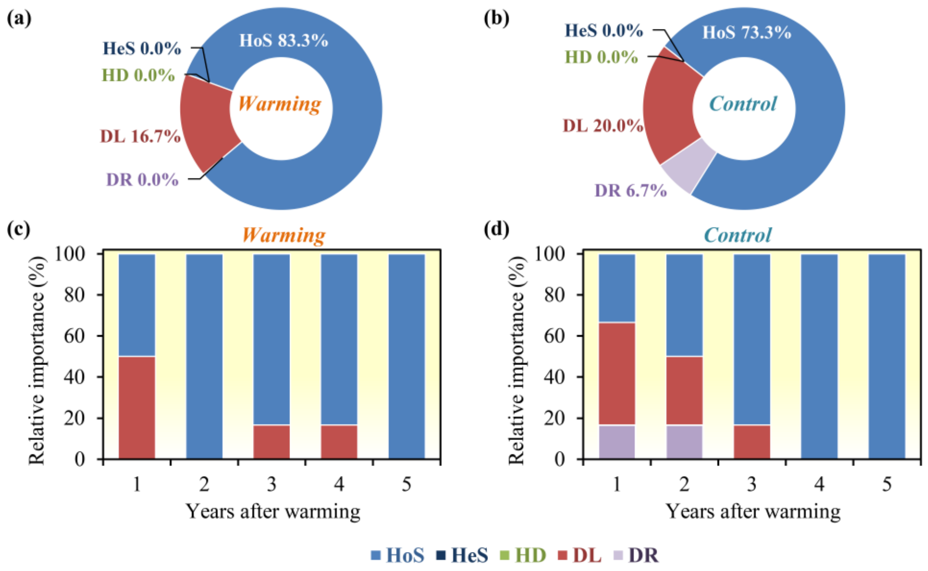


**Fig. S11. Relative importance of different assembly processes estimated with QPEN. (a)** Average relative importance of different processes in bacterial assembly under warming, and **(b)** under control. **(c)** Relative importance of different ecological processes under warming and **(d)** under control each year.

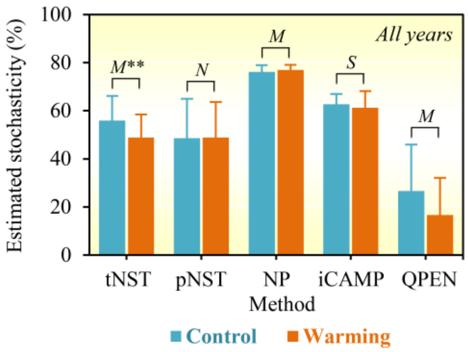


**Fig. S12. Stochasticity estimated by different methods with the experimental data across all 5 years after warming.** tNST and pNST represented taxonomic and phylogenetic normalized stochasticity ratio, respectively; NP represented abundance-weighted neutral taxa percentage. Significance was indicated as ***, $p < 0.01$; **, $p < 0.05$; L, M, S, and N represent large, medium, small, and negligible effect sizes, according to Cohen's *d* estimated as the mean difference between warming and control divided by pooled standard deviation. Error bars represented standard deviations.

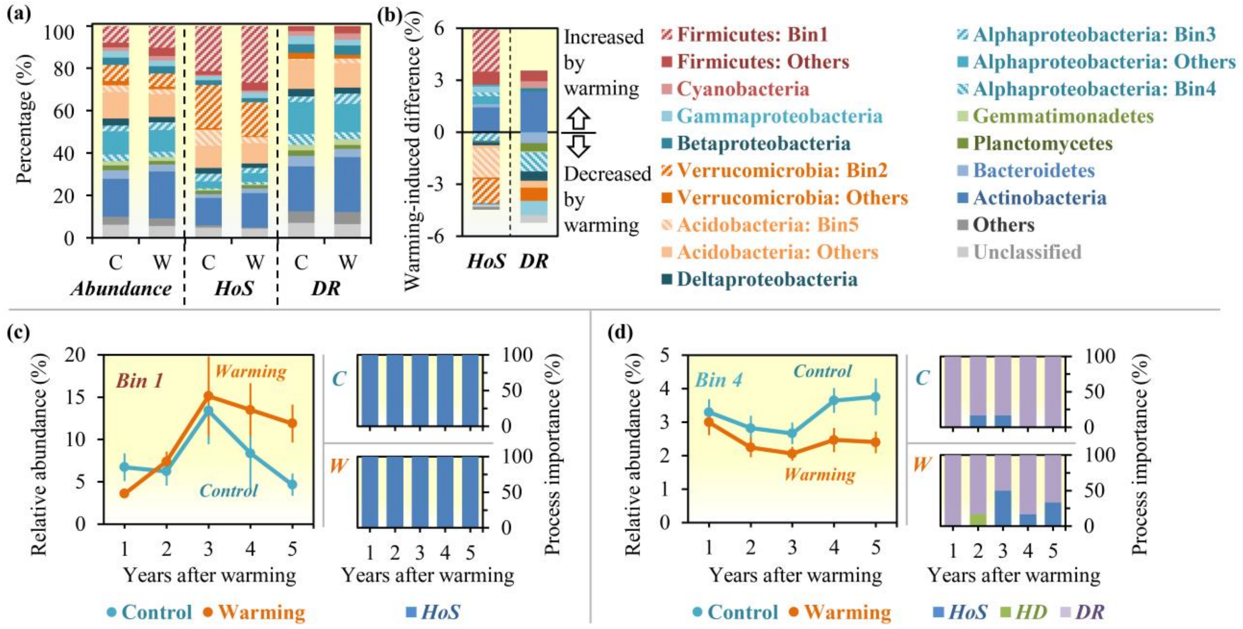


**Fig. S13. Ecological processes controlling major phylogenetic bins. (a)** Relative abundances of different phyla and their relative contributions to HoS and DR under control (C) and warming (W). Top 5 abundant bins were particularly highlighted. **(b)** Warming-induced changes of different phyla in the later 3 years. **(c)** and **(d),** relative abundances of Bin 1 and Bin 4, and relative importance of ecological processes controlling their assembly under control (C, aqua) and warming (W, orange). Error bars indicated standard error (n = 4).

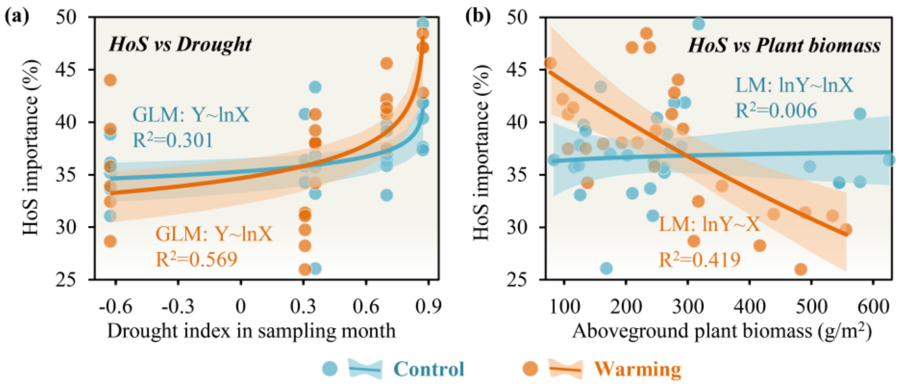


**Fig. S14. Correlation between relative importance of HoS and environmental variables under warming (orange) and control (aqua) (a)** HoS importance vs different degrees of drought in the sampling month. **(b)** HoS importance vs aboveground plant biomass. The shadow indicated 95% confidence interval; LM, linear model was used for Mantel test; GLM, general linear model was used for Mantel test; lnX or lnY, the values were log-transformed before fitting the model.

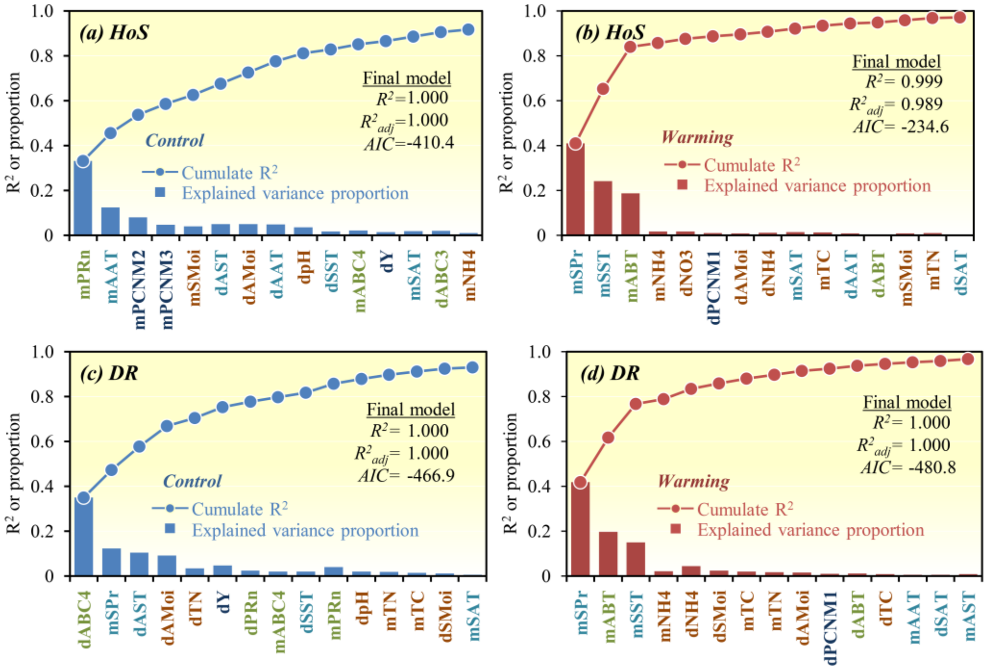


**Fig. S15. Multiple regression to link the importance of major processes with measured geographic, plant, edaphic, and climate factors by MRM analysis. (a)** MRM analysis of HoS under control **(b)** HoS under warming; **(c)** DR under control; **(d)** DR under warming. Factors were ranked as forward selected based on adjusted R square, of which the cumulated $R^2$ and variations, which were additionally explained by each factor were showed in the charts. Only the first 15 factors were listed. In the factor abbreviations, the first letter ("d" or "m") indicated whether the difference or the mean of the factor in each pair of samples were analyzed for correlation. ABC3, aboveground biomass of C-3 plants; ABC4, aboveground biomass of C-4 plants; ABT, aboveground biomass of all plants; PRn, plant richness; pH, soil pH; TC, soil total carbon; TN, soil total nitrogen; NO3, soil nitrate nitrogen; NH4, soil ammonium nitrogen; SMoi, soil moisture in sampling month; AMoi, annual mean of soil moisture; SPr, precipitation in sampling month; APr, annual precipitation; AST, annual mean of soil temperature; SST, mean soil temperature in the sampling month; AAT, annual mean of air temperature; SAT, mean air temperature in the sampling month; X and Y, geographic Cartesian coordinates, eastward and northward, respectively; Dist, geographic distance; PCNM1-5, principal coordinates of neighborhood matrix calculated from geographic distance.

#### Supplementary Tables

**Table S1. Defined (expected) relative importance of different ecological processes under various simulated situations.** Each simulated scenario has following 15 situations. The location of plot HA, HB, LA, and LB is showed in Fig. S2a. HeS, heterogeneous selection; HoS, homogeneous selection; DL, dispersal limitation; HD, homogenizing dispersal; DR, "drift", including drift, stochastic diversification, weak selection, and/or weak dispersal; ST, expected stochasticity.

| Situation | Selection | Dispersal | Drift | Comparison | HeS | HoS | DL | HD | DR | ST |
|---|---|---|---|---|---|---|---|---|---|---|
| **1** | 100% | 0% | 0% | HA vs HB | 0% | 100% | 0% | 0% | 0% | 0% |
| | | | | HA vs LA | 100% | 0% | 0% | 0% | 0% | |
| | | | | HA vs LB | 100% | 0% | 0% | 0% | 0% | |
| | | | | HB vs LA | 100% | 0% | 0% | 0% | 0% | |
| | | | | HB vs LB | 100% | 0% | 0% | 0% | 0% | |
| | | | | LA vs LB | 0% | 100% | 0% | 0% | 0% | |
| **2** | 0% | 100% | 0% | HA vs HB | 0% | 0% | 100% | 0% | 0% | 100% |
| | | | | HA vs LA | 0% | 0% | 0% | 100% | 0% | |
| | | | | HA vs LB | 0% | 0% | 100% | 0% | 0% | |
| | | | | HB vs LA | 0% | 0% | 100% | 0% | 0% | |
| | | | | HB vs LB | 0% | 0% | 0% | 100% | 0% | |
| | | | | LA vs LB | 0% | 0% | 100% | 0% | 50% | |
| **3** | 0% | 0% | 100% | HA vs HB | 0% | 0% | 0% | 0% | 100% | 100% |
| | | | | HA vs LA | 0% | 0% | 0% | 0% | 100% | |
| | | | | HA vs LB | 0% | 0% | 0% | 0% | 100% | |
| | | | | HB vs LA | 0% | 0% | 0% | 0% | 100% | |
| | | | | HB vs LB | 0% | 0% | 0% | 0% | 100% | |
| | | | | LA vs LB | 0% | 0% | 0% | 0% | 100% | |
| **4** | 0% | 25% | 75% | HA vs HB | 0% | 0% | 25% | 0% | 75% | 100% |
| | | | | HA vs LA | 0% | 0% | 0% | 25% | 75% | |
| | | | | HA vs LB | 0% | 0% | 25% | 0% | 75% | |
| | | | | HB vs LA | 0% | 0% | 25% | 0% | 75% | |
| | | | | HB vs LB | 0% | 0% | 0% | 25% | 75% | |
| | | | | LA vs LB | 0% | 0% | 25% | 0% | 75% | |
| **5** | 0% | 50% | 50% | HA vs HB | 0% | 0% | 50% | 0% | 50% | 100% |
| | | | | HA vs LA | 0% | 0% | 0% | 50% | 50% | |
| | | | | HA vs LB | 0% | 0% | 50% | 0% | 50% | |
| | | | | HB vs LA | 0% | 0% | 50% | 0% | 50% | |
| | | | | HB vs LB | 0% | 0% | 0% | 50% | 50% | |
| | | | | LA vs LB | 0% | 0% | 50% | 0% | 50% | |
| **6** | 0% | 75% | 25% | HA vs HB | 0% | 0% | 75% | 0% | 25% | 100% |
| | | | | HA vs LA | 0% | 0% | 0% | 75% | 25% | |
| | | | | HA vs LB | 0% | 0% | 75% | 0% | 25% | |
| | | | | HB vs LA | 0% | 0% | 75% | 0% | 25% | |
| | | | | HB vs LB | 0% | 0% | 0% | 75% | 25% | |
| | | | | LA vs LB | 0% | 0% | 75% | 0% | 25% | |
| **7** | 25% | 0% | 75% | HA vs HB | 0% | 25% | 0% | 0% | 75% | 75% |
| | | | | HA vs LA | 25% | 0% | 0% | 0% | 75% | |
| | | | | HA vs LB | 25% | 0% | 0% | 0% | 75% | |
| | | | | HB vs LA | 25% | 0% | 0% | 0% | 75% | |
| | | | | HB vs LB | 25% | 0% | 0% | 0% | 75% | |
| | | | | LA vs LB | 0% | 25% | 0% | 0% | 75% | |

**Table S1. Continued**

| Situation | Selection | Dispersal | Drift | Comparison | HeS | HoS | DL | HD | DR | ST |
|---|---|---|---|---|---|---|---|---|---|---|
| **8** | 50% | 0% | 50% | HA vs HB | 0% | 50% | 0% | 0% | 50% | 50% |
| | | | | HA vs LA | 50% | 0% | 0% | 0% | 50% | |
| | | | | HA vs LB | 50% | 0% | 0% | 0% | 50% | |
| | | | | HB vs LA | 50% | 0% | 0% | 0% | 50% | |
| | | | | HB vs LB | 50% | 0% | 0% | 0% | 50% | |
| | | | | LA vs LB | 0% | 50% | 0% | 0% | 50% | |
| **9** | 75% | 0% | 25% | HA vs HB | 0% | 75% | 0% | 0% | 25% | 25% |
| | | | | HA vs LA | 75% | 0% | 0% | 0% | 25% | |
| | | | | HA vs LB | 75% | 0% | 0% | 0% | 25% | |
| | | | | HB vs LA | 75% | 0% | 0% | 0% | 25% | |
| | | | | HB vs LB | 75% | 0% | 0% | 0% | 25% | |
| | | | | LA vs LB | 0% | 75% | 0% | 0% | 25% | |
| **10** | 25% | 75% | 0% | HA vs HB | 0% | 25% | 75% | 0% | 0% | 75% |
| | | | | HA vs LA | 25% | 0% | 0% | 75% | 0% | |
| | | | | HA vs LB | 25% | 0% | 75% | 0% | 0% | |
| | | | | HB vs LA | 25% | 0% | 75% | 0% | 0% | |
| | | | | HB vs LB | 25% | 0% | 0% | 75% | 0% | |
| | | | | LA vs LB | 0% | 25% | 75% | 0% | 0% | |
| **11** | 50% | 50% | 0% | HA vs HB | 0% | 50% | 50% | 0% | 0% | 50% |
| | | | | HA vs LA | 50% | 0% | 0% | 50% | 0% | |
| | | | | HA vs LB | 50% | 0% | 50% | 0% | 0% | |
| | | | | HB vs LA | 50% | 0% | 50% | 0% | 0% | |
| | | | | HB vs LB | 50% | 0% | 0% | 50% | 0% | |
| | | | | LA vs LB | 0% | 50% | 50% | 0% | 0% | |
| **12** | 75% | 25% | 0% | HA vs HB | 0% | 75% | 25% | 0% | 0% | 25% |
| | | | | HA vs LA | 75% | 0% | 0% | 25% | 0% | |
| | | | | HA vs LB | 75% | 0% | 25% | 0% | 0% | |
| | | | | HB vs LA | 75% | 0% | 25% | 0% | 0% | |
| | | | | HB vs LB | 75% | 0% | 0% | 25% | 0% | |
| | | | | LA vs LB | 0% | 75% | 25% | 0% | 0% | |
| **13** | 25% | 25% | 50% | HA vs HB | 0% | 25% | 25% | 0% | 50% | 75% |
| | | | | HA vs LA | 25% | 0% | 0% | 25% | 50% | |
| | | | | HA vs LB | 25% | 0% | 25% | 0% | 50% | |
| | | | | HB vs LA | 25% | 0% | 25% | 0% | 50% | |
| | | | | HB vs LB | 25% | 0% | 0% | 25% | 50% | |
| | | | | LA vs LB | 0% | 25% | 25% | 0% | 50% | |
| **14** | 25% | 50% | 25% | HA vs HB | 0% | 25% | 50% | 0% | 25% | 75% |
| | | | | HA vs LA | 25% | 0% | 0% | 50% | 25% | |
| | | | | HA vs LB | 25% | 0% | 50% | 0% | 25% | |
| | | | | HB vs LA | 25% | 0% | 50% | 0% | 25% | |
| | | | | HB vs LB | 25% | 0% | 0% | 50% | 25% | |
| | | | | LA vs LB | 0% | 25% | 50% | 0% | 25% | |
| **15** | 50% | 25% | 25% | HA vs HB | 0% | 50% | 25% | 0% | 25% | 50% |
| | | | | HA vs LA | 50% | 0% | 0% | 25% | 25% | |
| | | | | HA vs LB | 50% | 0% | 25% | 0% | 25% | |
| | | | | HB vs LA | 50% | 0% | 25% | 0% | 25% | |
| | | | | HB vs LB | 50% | 0% | 0% | 25% | 25% | |
| | | | | LA vs LB | 0% | 50% | 25% | 0% | 25% | |

**Table S2. Coefficient of determination ($R^2$) between the relative importance of HoS or DR and each factor.** The relative importance of HoS and DR was estimated by iCAMP. The correlation was analyzed by modified Mantel test based on linear model (LM) and general linear model (GLM). The relative importance of an ecological process (Y) and each factor (X) were either log-transformed or not before fitting the models, to test linear (Y ~ X), logarithmic (Y ~ lnX), exponential (lnY ~ X), and power law (lnY ~ lnX) relationship. The best model was selected based on $R^2$. In the factor abbreviations, the first letter ("d" or "m") indicated whether the difference or the mean of the factor for each pair of samples were performed by correlation analysis; ABC3, aboveground biomass of C-3 plants; ABC4, aboveground biomass of C-4 plants; ABT, aboveground biomass of all plants; PRn, plant richness; pH, soil pH; TC, total soil carbon; TN, total soil nitrogen; NO3, soil nitrate ; NH4, soil ammonium nitrogen; SMoi, soil moisture in sampling month; AMoi, annual mean of soil moisture; SPr, precipitation in sampling month; APr, annual precipitation; SDI, drought index in the sampling month, which is additive inverse of the standardized precipitation-evapotranspiration index (SPEI) in the sampling month; ADI, annual mean of the drought index, which was additive inverse of annual mean SPEI; AST, annual mean of soil temperature; SST, mean soil temperature in sampling month; AAT, annual mean of air temperature; SAT, mean air temperature in sampling month; WY, warming years; X, eastward coordinate; Y, northward coordinate; Dist, geographic distance; PCNM1-5, principal coordinates of neighborhood matrix calculated from geographic distance. Significance is indicated as ***, $p < 0.01$; **, $p < 0.05$; *, $p < 0.1$.

| X: factor | Control | | | | | | Warming | | | | | | Control | Warming |
|---|---|---|---|---|---|---|---|---|---|---|---|---|---|---|
| | Mantel-LM | | | | Mantel-GLM | | Mantel-LM | | | | Mantel-GLM | | Best model results | |
| | Y~X | Y~lnX | lnY~X | lnY~lnX | Y~X | Y~lnX | Y~X | Y~lnX | lnY~X | lnY~lnX | Y~X | Y~lnX | | |
| **(1) Y: HoS importance** | | | | | | | | | | | | | | |
| **dABC3** | 0.002 | 0.013 | 0.000 | 0.007 | 0.002 | 0.013 | 0.007 | 0.004 | 0.008 | 0.009 | 0.007 | 0.004 | 0.013 | 0.009 |
| **mABC3** | 0.000 | 0.001 | 0.000 | 0.000 | 0.000 | 0.001 | 0.138 | 0.063 | 0.181 | 0.089 | 0.135 | 0.062 | 0.001 | 0.181 |
| **dABC4** | 0.241** | 0.345*** | 0.218** | 0.304*** | 0.244** | 0.347*** | 0.065 | 0.049 | 0.048 | 0.033 | 0.067 | 0.051 | 0.347*** | 0.067 |
| **mABC4** | 0.003 | 0.046 | 0.005 | 0.038 | 0.003 | 0.045 | 0.328* | 0.498* | 0.292* | 0.441* | 0.338* | 0.500* | 0.046 | 0.500* |
| **dABT** | 0.118* | 0.103* | 0.129* | 0.120* | 0.116* | 0.102* | 0.042 | 0.012 | 0.037 | 0.022 | 0.042 | 0.012 | 0.129* | 0.042 |
| **mABT** | 0.000 | 0.006 | 0.000 | 0.006 | 0.000 | 0.005 | 0.370* | 0.255 | 0.419* | 0.297 | 0.364* | 0.248 | 0.006 | 0.419* |
| **dPRn** | 0.003 | 0.010 | 0.003 | 0.010 | 0.003 | 0.010 | 0.101* | 0.128 | 0.102* | 0.131 | 0.100* | 0.126 | 0.010 | 0.131 |
| **mPRn** | 0.332*** | 0.278*** | 0.323*** | 0.271** | 0.336*** | 0.277*** | 0.016 | 0.005 | 0.016 | 0.006 | 0.016 | 0.005 | 0.336*** | 0.016 |
| **dpH** | 0.001 | 0.001 | 0.001 | 0.001 | 0.001 | 0.001 | 0.087 | 0.083 | 0.078 | 0.074 | 0.087 | 0.083 | 0.001 | 0.087 |
| **mpH** | 0.022 | 0.022 | 0.027 | 0.027 | 0.022 | 0.022 | 0.109 | 0.103 | 0.129 | 0.123 | 0.107 | 0.101 | 0.027 | 0.129 |
| **dTC** | 0.004 | 0.009 | 0.002 | 0.007 | 0.004 | 0.009 | 0.000 | 0.000 | 0.002 | 0.004 | 0.000 | 0.000 | 0.009 | 0.004 |
| **mTC** | 0.007 | 0.009 | 0.007 | 0.007 | 0.007 | 0.009 | 0.028 | 0.031 | 0.032 | 0.032 | 0.028 | 0.031 | 0.009 | 0.032 |
| **dTN** | 0.012 | 0.022 | 0.009 | 0.018 | 0.012 | 0.022 | 0.001 | 0.000 | 0.000 | 0.002 | 0.001 | 0.000 | 0.022 | 0.002 |
| **mTN** | 0.013 | 0.018 | 0.012 | 0.016 | 0.013 | 0.018 | 0.011 | 0.013 | 0.012 | 0.012 | 0.011 | 0.013 | 0.018 | 0.013 |
| **dNO3** | 0.006 | 0.019 | 0.007 | 0.021 | 0.006 | 0.019 | 0.049 | 0.223* | 0.046 | 0.207* | 0.049 | 0.223* | 0.021 | 0.223* |
| **mNO3** | 0.011 | 0.067 | 0.012 | 0.069 | 0.011 | 0.067 | 0.001 | 0.046 | 0.001 | 0.056 | 0.001 | 0.044 | 0.069 | 0.056 |
| **dNH4** | 0.053 | 0.034 | 0.061 | 0.040 | 0.053 | 0.034 | 0.022 | 0.045 | 0.019 | 0.041 | 0.022 | 0.044 | 0.061 | 0.045 |
| **mNH4** | 0.134* | 0.152* | 0.142* | 0.158* | 0.133* | 0.152* | 0.000 | 0.002 | 0.000 | 0.002 | 0.000 | 0.002 | 0.158* | 0.002 |

**Table S2. Continued**

| X: factor | Control | | | | | | Warming | | | | | | Control | Warming |
|---|---|---|---|---|---|---|---|---|---|---|---|---|---|---|
| | Mantel-LM | | | | Mantel-GLM | | Mantel-LM | | | | Mantel-GLM | | Best model results | |
| | Y~X | Y~lnX | lnY~X | lnY~lnX | Y~X | Y~lnX | Y~X | Y~lnX | lnY~X | lnY~lnX | Y~X | Y~lnX | | |
| **(1) Y: HoS importance** | | | | | | | | | | | | | | |
| **dSMoi** | 0.002 | 0.002 | 0.001 | 0.005 | 0.002 | 0.002 | 0.102 | 0.011 | 0.131 | 0.021 | 0.099 | 0.011 | 0.005 | 0.131 |
| **mSMoi** | 0.003 | 0.005 | 0.002 | 0.004 | 0.003 | 0.005 | 0.046 | 0.014 | 0.073 | 0.027 | 0.045 | 0.014 | 0.005 | 0.073 |
| **dAMoi** | 0.228** | 0.273*** | 0.264*** | 0.310*** | 0.226** | 0.271*** | 0.006 | 0.009 | 0.006 | 0.013 | 0.006 | 0.009 | 0.310*** | 0.013 |
| **mAMoi** | 0.011 | 0.011 | 0.007 | 0.008 | 0.011 | 0.011 | 0.080 | 0.058 | 0.109 | 0.081 | 0.078 | 0.056 | 0.011 | 0.109 |
| **mSPr** | 0.253*** | 0.305*** | 0.234*** | 0.281*** | 0.256*** | 0.305*** | 0.411* | 0.526* | 0.367 | 0.464* | 0.423* | 0.527* | 0.305*** | 0.527* |
| **mAPr** | 0.009 | 0.004 | 0.013 | 0.007 | 0.009 | 0.004 | 0.083 | 0.068 | 0.084 | 0.071 | 0.082 | 0.067 | 0.013 | 0.084 |
| **mSDI** | 0.149 | 0.300** | 0.138 | 0.278** | 0.151 | 0.301** | 0.254 | 0.567* | 0.235 | 0.511* | 0.264 | 0.569* | 0.301** | 0.569* |
| **mADI** | 0.008 | 0.013 | 0.011 | 0.010 | 0.008 | 0.013 | 0.028 | 0.003 | 0.023 | 0.003 | 0.027 | 0.003 | 0.013 | 0.028 |
| **dAST** | 0.031 | 0.029 | 0.027 | 0.025 | 0.031 | 0.029 | 0.001 | 0.001 | 0.001 | 0.000 | 0.001 | 0.001 | 0.031 | 0.001 |
| **mAST** | 0.008 | 0.010 | 0.007 | 0.008 | 0.008 | 0.009 | 0.217 | 0.224 | 0.204 | 0.209 | 0.216 | 0.222 | 0.010 | 0.224 |
| **dSST** | 0.006 | 0.004 | 0.007 | 0.005 | 0.006 | 0.004 | 0.077 | 0.135 | 0.085 | 0.147* | 0.078 | 0.136 | 0.007 | 0.147* |
| **mSST** | 0.005 | 0.004 | 0.007 | 0.006 | 0.005 | 0.004 | 0.313* | 0.321* | 0.314* | 0.324* | 0.313* | 0.322* | 0.007 | 0.324* |
| **dAAT** | 0.037 | 0.041 | 0.034 | 0.039 | 0.037 | 0.042 | 0.067 | 0.042 | 0.074 | 0.050 | 0.067 | 0.041 | 0.042 | 0.074 |
| **mAAT** | 0.043 | 0.041 | 0.041 | 0.039 | 0.043 | 0.042 | 0.056 | 0.047 | 0.036 | 0.029 | 0.057 | 0.049 | 0.043 | 0.057 |
| **dSAT** | 0.050 | 0.045 | 0.059 | 0.053 | 0.050 | 0.045 | 0.021 | 0.019 | 0.031 | 0.029 | 0.020 | 0.019 | 0.059 | 0.031 |
| **mSAT** | 0.040 | 0.035 | 0.048 | 0.043 | 0.039 | 0.035 | 0.043 | 0.043 | 0.038 | 0.038 | 0.044 | 0.043 | 0.048 | 0.044 |
| **mWY** | | | | | | | 0.262 | 0.376 | 0.312 | 0.434 | 0.254 | 0.368 | | 0.434 |
| **dX** | 0.008 | 0.006 | 0.011 | 0.010 | 0.008 | 0.006 | 0.000 | 0.000 | 0.000 | 0.000 | 0.000 | 0.000 | 0.011 | 0.000 |
| **mX** | 0.010 | 0.000 | 0.008 | 0.000 | 0.010 | 0.000 | 0.000 | 0.000 | 0.000 | 0.000 | 0.000 | 0.000 | 0.010 | 0.000 |
| **dY** | 0.002 | 0.002 | 0.005 | 0.005 | 0.002 | 0.002 | 0.006 | 0.000 | 0.006 | 0.000 | 0.006 | 0.000 | 0.005 | 0.006 |
| **mY** | 0.000 | 0.000 | 0.000 | 0.000 | 0.000 | 0.000 | 0.008* | 0.007** | 0.010** | 0.011*** | 0.008* | 0.008** | 0.000 | 0.011*** |
| **Dist** | 0.001 | 0.000 | 0.000 | 0.000 | 0.001 | 0.000 | 0.005 | 0.005 | 0.005 | 0.004 | 0.005 | 0.005 | 0.001 | 0.005 |
| **dPCNM1** | 0.000 | 0.005 | 0.003 | 0.007 | 0.000 | 0.005 | 0.007 | 0.011 | 0.008 | 0.014 | 0.007 | 0.011 | 0.007 | 0.014 |
| **mPCNM1** | 0.003 | 0.005 | 0.003 | 0.006 | 0.003 | 0.005 | 0.008** | 0.008* | 0.010*** | 0.009** | 0.008** | 0.008* | 0.006 | 0.010*** |
| **dPCNM2** | 0.001 | 0.035 | 0.001 | 0.034 | 0.001 | 0.035 | 0.000 | 0.001 | 0.000 | 0.001 | 0.000 | 0.001 | 0.035 | 0.001 |
| **mPCNM2** | 0.016 | 0.036* | 0.014 | 0.035* | 0.016 | 0.036* | 0.000 | 0.001 | 0.000 | 0.001 | 0.000 | 0.001 | 0.036* | 0.001 |
| **dPCNM3** | 0.001 | 0.006 | 0.002 | 0.009 | 0.001 | 0.006 | 0.002 | 0.001 | 0.002 | 0.001 | 0.002 | 0.001 | 0.009 | 0.002 |
| **mPCNM3** | 0.045* | 0.007 | 0.051* | 0.010 | 0.044* | 0.007 | 0.000 | 0.000 | 0.001 | 0.000 | 0.000 | 0.000 | 0.051* | 0.001 |
| **dPCNM4** | 0.012 | 0.008 | 0.012 | 0.010 | 0.012 | 0.008 | 0.002 | 0.000 | 0.003 | 0.000 | 0.002 | 0.000 | 0.012 | 0.003 |
| **mPCNM4** | 0.000 | 0.002 | 0.000 | 0.003 | 0.000 | 0.002 | 0.002 | 0.000 | 0.002 | 0.000 | 0.002 | 0.000 | 0.003 | 0.002 |
| **dPCNM5** | 0.016 | 0.011 | 0.021 | 0.015 | 0.016 | 0.011 | 0.004 | 0.008* | 0.003 | 0.010* | 0.004 | 0.008* | 0.021 | 0.010* |
| **mPCNM5** | 0.001 | 0.004 | 0.003 | 0.006 | 0.001 | 0.004 | 0.001 | 0.007* | 0.001 | 0.009 | 0.001 | 0.007* | 0.006 | 0.009 |

**Table S2. Continued**

| X: factor | Control | | | | | | Warming | | | | | | Control | Warming |
|---|---|---|---|---|---|---|---|---|---|---|---|---|---|---|
| | Mantel-LM | | | | Mantel-GLM | | Mantel-LM | | | | Mantel-GLM | | Best model results | |
| | Y~X | Y~lnX | lnY~X | lnY~lnX | Y~X | Y~lnX | Y~X | Y~lnX | lnY~X | lnY~lnX | Y~X | Y~lnX | | |
| **(2) Y: DR importance** | | | | | | | | | | | | | | |
| **dABC3** | 0.088 | 0.036 | 0.093 | 0.037 | 0.087 | 0.036 | 0.008 | 0.008 | 0.007 | 0.005 | 0.008 | 0.008 | 0.093 | 0.008 |
| **mABC3** | 0.016 | 0.020 | 0.014 | 0.020 | 0.016 | 0.020 | 0.130 | 0.067 | 0.108 | 0.054 | 0.128 | 0.067 | 0.020 | 0.130 |
| **dABC4** | 0.350*** | 0.442*** | 0.364*** | 0.469*** | 0.354*** | 0.443*** | 0.094 | 0.058 | 0.109 | 0.072 | 0.096 | 0.059 | 0.469*** | 0.109 |
| **mABC4** | 0.021 | 0.039 | 0.019 | 0.041 | 0.021 | 0.039 | 0.339* | 0.481* | 0.356* | 0.507* | 0.347* | 0.482* | 0.041 | 0.507* |
| **dABT** | 0.024 | 0.078 | 0.026 | 0.074 | 0.024 | 0.077 | 0.048 | 0.018 | 0.051 | 0.012 | 0.049 | 0.018 | 0.078 | 0.051 |
| **mABT** | 0.025 | 0.046 | 0.022 | 0.044 | 0.024 | 0.046 | 0.361* | 0.278 | 0.332 | 0.253 | 0.358* | 0.274 | 0.046 | 0.361* |
| **dPRn** | 0.001 | 0.018 | 0.001 | 0.017 | 0.001 | 0.018 | 0.077 | 0.087 | 0.076 | 0.087 | 0.077 | 0.087 | 0.018 | 0.087 |
| **mPRn** | 0.329*** | 0.290** | 0.323*** | 0.284** | 0.332*** | 0.290** | 0.011 | 0.008 | 0.010 | 0.007 | 0.011 | 0.007 | 0.332*** | 0.011 |
| **dpH** | 0.000 | 0.001 | 0.000 | 0.000 | 0.000 | 0.001 | 0.103 | 0.099 | 0.103* | 0.099 | 0.103 | 0.099 | 0.001 | 0.103 |
| **mpH** | 0.000 | 0.000 | 0.000 | 0.000 | 0.000 | 0.000 | 0.098 | 0.093 | 0.086 | 0.081 | 0.097 | 0.092 | 0.000 | 0.098 |
| **dTC** | 0.036 | 0.048 | 0.040 | 0.053 | 0.037 | 0.048 | 0.001 | 0.001 | 0.002 | 0.002 | 0.001 | 0.001 | 0.053 | 0.002 |
| **mTC** | 0.000 | 0.003 | 0.000 | 0.003 | 0.000 | 0.003 | 0.037 | 0.046 | 0.031 | 0.041 | 0.037 | 0.046 | 0.003 | 0.046 |
| **dTN** | 0.068 | 0.087 | 0.069 | 0.089* | 0.068 | 0.087 | 0.003 | 0.001 | 0.004 | 0.002 | 0.003 | 0.001 | 0.089* | 0.004 |
| **mTN** | 0.001 | 0.003 | 0.001 | 0.003 | 0.001 | 0.003 | 0.014 | 0.019 | 0.012 | 0.017 | 0.015 | 0.020 | 0.003 | 0.020 |
| **dNO3** | 0.025 | 0.009 | 0.019 | 0.008 | 0.025 | 0.009 | 0.039 | 0.137* | 0.038 | 0.140* | 0.038 | 0.137* | 0.025 | 0.140* |
| **mNO3** | 0.001 | 0.010 | 0.000 | 0.013 | 0.001 | 0.010 | 0.003 | 0.024 | 0.003 | 0.022 | 0.003 | 0.024 | 0.013 | 0.024 |
| **dNH4** | 0.032 | 0.014 | 0.027 | 0.011 | 0.032 | 0.014 | 0.042 | 0.093 | 0.041 | 0.088 | 0.042 | 0.093 | 0.032 | 0.093 |
| **mNH4** | 0.123 | 0.157* | 0.111 | 0.145* | 0.122 | 0.156* | 0.000 | 0.002 | 0.000 | 0.002 | 0.000 | 0.002 | 0.157* | 0.002 |
| **dSMoi** | 0.008 | 0.001 | 0.008 | 0.000 | 0.008 | 0.001 | 0.068 | 0.004 | 0.053 | 0.001 | 0.067 | 0.004 | 0.008 | 0.068 |
| **mSMoi** | 0.000 | 0.001 | 0.000 | 0.001 | 0.000 | 0.001 | 0.049 | 0.010 | 0.035 | 0.005 | 0.049 | 0.010 | 0.001 | 0.049 |
| **dAMoi** | 0.116* | 0.153* | 0.107* | 0.142* | 0.115* | 0.152* | 0.002 | 0.005 | 0.001 | 0.003 | 0.002 | 0.005 | 0.153* | 0.005 |
| **mAMoi** | 0.039 | 0.039 | 0.039 | 0.040 | 0.039 | 0.039 | 0.077 | 0.055 | 0.062 | 0.044 | 0.076 | 0.055 | 0.040 | 0.077 |
| **mSPr** | 0.333** | 0.355** | 0.332** | 0.362** | 0.335** | 0.354** | 0.419* | 0.493* | 0.435* | 0.520* | 0.426* | 0.492* | 0.362** | 0.520* |
| **mAPr** | 0.003 | 0.008 | 0.002 | 0.007 | 0.002 | 0.008 | 0.049 | 0.037 | 0.050 | 0.037 | 0.048 | 0.036 | 0.008 | 0.050 |
| **mSDI** | 0.221 | 0.341** | 0.216 | 0.347** | 0.223 | 0.342** | 0.284 | 0.539* | 0.288 | 0.561** | 0.291 | 0.539* | 0.347** | 0.561** |
| **mADI** | 0.001 | 0.062 | 0.001 | 0.056 | 0.001 | 0.061 | 0.009 | 0.016 | 0.012 | 0.015 | 0.009 | 0.016 | 0.062 | 0.016 |
| **dAST** | 0.056 | 0.051 | 0.056 | 0.051 | 0.056 | 0.051 | 0.001 | 0.004 | 0.001 | 0.004 | 0.001 | 0.004 | 0.056 | 0.004 |
| **mAST** | 0.000 | 0.000 | 0.000 | 0.000 | 0.000 | 0.000 | 0.134 | 0.137 | 0.140 | 0.144 | 0.134 | 0.137 | 0.000 | 0.144 |
| **dSST** | 0.001 | 0.000 | 0.001 | 0.000 | 0.001 | 0.000 | 0.123 | 0.184* | 0.115 | 0.174* | 0.124 | 0.186* | 0.001 | 0.186* |
| **mSST** | 0.014 | 0.011 | 0.011 | 0.009 | 0.014 | 0.011 | 0.227 | 0.237 | 0.224 | 0.233 | 0.227 | 0.237 | 0.014 | 0.237 |
| **dAAT** | 0.074 | 0.079 | 0.074 | 0.079 | 0.074 | 0.079 | 0.132* | 0.100 | 0.128* | 0.093 | 0.132* | 0.099 | 0.079 | 0.132* |
| **mAAT** | 0.029 | 0.029 | 0.033 | 0.033 | 0.029 | 0.029 | 0.030 | 0.023 | 0.041 | 0.033 | 0.031 | 0.024 | 0.033 | 0.041 |
| **dSAT** | 0.023 | 0.020 | 0.021 | 0.018 | 0.023 | 0.020 | 0.000 | 0.000 | 0.000 | 0.000 | 0.000 | 0.000 | 0.023 | 0.000 |

**Table S2. Continued**

| X: factor | Control | | | | | | Warming | | | | | | Control | Warming |
|---|---|---|---|---|---|---|---|---|---|---|---|---|---|---|
| | Mantel-LM | | | | Mantel-GLM | | Mantel-LM | | | | Mantel-GLM | | Best model results | |
| | Y~X | Y~lnX | lnY~X | lnY~lnX | Y~X | Y~lnX | Y~X | Y~lnX | lnY~X | lnY~lnX | Y~X | Y~lnX | | |
| **(2) Y: DR importance** | | | | | | | | | | | | | | |
| **mSAT** | 0.025 | 0.021 | 0.021 | 0.018 | 0.025 | 0.021 | 0.047 | 0.047 | 0.051 | 0.051 | 0.047 | 0.047 | 0.025 | 0.051 |
| **mWY** | | | | | | | 0.256 | 0.350 | 0.225 | 0.315 | 0.251 | 0.346 | | 0.350 |
| **dX** | 0.013 | 0.002 | 0.011 | 0.001 | 0.013 | 0.002 | 0.000 | 0.000 | 0.000 | 0.000 | 0.000 | 0.000 | 0.013 | 0.000 |
| **mX** | 0.031 | 0.010 | 0.036 | 0.013 | 0.031 | 0.010 | 0.001 | 0.001 | 0.000 | 0.000 | 0.001 | 0.001 | 0.036 | 0.001 |
| **dY** | 0.000 | 0.000 | 0.000 | 0.000 | 0.000 | 0.000 | 0.005 | 0.008 | 0.005 | 0.009 | 0.005 | 0.008 | 0.000 | 0.009 |
| **mY** | 0.001 | 0.001 | 0.001 | 0.001 | 0.001 | 0.001 | 0.002 | 0.002 | 0.003 | 0.004 | 0.002 | 0.002 | 0.001 | 0.004 |
| **Dist** | 0.004 | 0.002 | 0.006 | 0.004 | 0.004 | 0.002 | 0.002 | 0.002 | 0.002 | 0.003 | 0.002 | 0.002 | 0.006 | 0.003 |
| **dPCNM1** | 0.004 | 0.000 | 0.008 | 0.001 | 0.004 | 0.000 | 0.011 | 0.008 | 0.010 | 0.006 | 0.011 | 0.008 | 0.008 | 0.011 |
| **mPCNM1** | 0.000 | 0.000 | 0.000 | 0.001 | 0.000 | 0.000 | 0.001 | 0.001 | 0.002 | 0.001 | 0.001 | 0.001 | 0.001 | 0.002 |
| **dPCNM2** | 0.000 | 0.043 | 0.000 | 0.046 | 0.000 | 0.043 | 0.000 | 0.002 | 0.000 | 0.002 | 0.000 | 0.002 | 0.046 | 0.002 |
| **mPCNM2** | 0.046* | 0.045 | 0.050** | 0.048 | 0.046* | 0.045 | 0.000 | 0.002 | 0.000 | 0.002 | 0.000 | 0.002 | 0.050** | 0.002 |
| **dPCNM3** | 0.009 | 0.000 | 0.011 | 0.000 | 0.009 | 0.000 | 0.013 | 0.008 | 0.015** | 0.010 | 0.013 | 0.008 | 0.011 | 0.015** |
| **mPCNM3** | 0.037 | 0.000 | 0.036 | 0.000 | 0.037 | 0.000 | 0.008* | 0.008* | 0.009 | 0.008 | 0.008* | 0.008* | 0.037 | 0.009 |
| **dPCNM4** | 0.000 | 0.000 | 0.000 | 0.000 | 0.000 | 0.000 | 0.004 | 0.000 | 0.003 | 0.000 | 0.004 | 0.000 | 0.000 | 0.004 |
| **mPCNM4** | 0.011 | 0.004 | 0.013 | 0.005 | 0.011 | 0.004 | 0.004 | 0.000 | 0.003 | 0.000 | 0.004 | 0.000 | 0.013 | 0.004 |
| **dPCNM5** | 0.004 | 0.002 | 0.003 | 0.001 | 0.004 | 0.002 | 0.001 | 0.001 | 0.001 | 0.001 | 0.001 | 0.001 | 0.004 | 0.001 |
| **mPCNM5** | 0.001 | 0.000 | 0.002 | 0.000 | 0.001 | 0.000 | 0.003 | 0.001 | 0.002 | 0.001 | 0.003 | 0.001 | 0.002 | 0.003 |
